# Cytological and genetic consequences for the progeny of a mitotic catastrophe provoked by Topoisomerase II deficiency

**DOI:** 10.1101/668954

**Authors:** Cristina Ramos-Pérez, Margaret Dominska, Laura Anaissi-Afonso, Sara Cazorla-Rivero, Oliver Quevedo, Isabel Lorenzo-Castrillejo, Thomas D Petes, Félix Machín

## Abstract

Topoisomerase II (Top2) removes topological linkages between replicated chromosomes. Top2 inhibition leads to mitotic catastrophe (MC) when cells unsuccessfully try to split their genetic material between the two daughter cells. Herein, we have characterized the fate of these daughter cells in the budding yeast. Clonogenic and microcolony experiments, in combination with vital and apoptotic stains, showed that 75% of daughter cells become senescent in the short term; they are unable to divide but remain alive. Decline in cell vitality then occurred, yet slowly, uncoordinatedly when comparing pairs of daughters, and independently of the cell death mediator Mca1/Yca1. Furthermore, we showed that senescence can be modulated by ploidy, suggesting that gross chromosome imbalances during segregation may account for this phenotype. Indeed, we found that diploid long-term survivors of the MC are prone to genomic imbalances such as trisomies, uniparental disomies and terminal loss of heterozygosity (LOH), the latter affecting the longest chromosome arms.

## INTRODUCTION

Mitotic catastrophe (MC) is a class of cell death still poorly understood, and with a conflictive definition among the scientific community [1–3]. In its most general acceptation, we can consider MC as the cell-death-triggering event that follows an aberrant mitosis. MC is presumed to be of the utmost importance in cancer biology, both as an oncosuppressive barrier in carcinogenesis and as a mechanism of cell death after anti-cancer treatments. Many antitumor drugs that damage the DNA or the microtubules lead to chromosome segregation failures, provided that cells do not stop their division cycle in a timely fashion [4, 5]. Human cells make use of checkpoints to arrest the cell cycle in G_1_/S or G_2_/M following treatment with these antitumor drugs, and tumour cells frequently lack one or several of these checkpoints. When checkpoints are functional, cancer cells often die after a transient cell cycle arrest through a regulated cell death (RCD) known as intrinsic apoptosis [6]. Apoptosis causes the permeabilization of the mitochondrial outer membrane and the subsequent leakage of pro-apoptotic factors into the cytosol. Execution of the intrinsic apoptosis is significatively accelerated by the activation of the so-called caspase-mediated transduction cascade. When checkpoints or apoptosis are non-functional, MC is expected to ensue. Therefore, understanding MC is becoming increasingly important in cancer biology.

MC is expected to kill most of the progeny due to major genomic imbalances and massive irreparable DNA damage. However, whether MC might also trigger a form of RCD or accidental cell death (ACD) is still unclear. In addition, MC is reported to lead to senescence in certain backgrounds [1, 7]. Senescence refers to the irreversible cell cycle arrest of otherwise live cells. Significant differences in the outcome are expected for MCs that result from different sources. Thus, MC occurring upon microtubule damage likely leads to missegregation of whole chromosomes prior to cell death, whereas DNA damage can give rise to more complex outcomes. For instance, DNA damage may result in either breakage of the DNA molecule (i.e, double strand breaks, DSBs) or replication stress (for example, stalled replication forks). Attempts to segregate broken chromosomes would yield daughter cells with irreparable damage. Attempts to segregate underreplicated chromosomes would cause the formation of anaphase bridges, which could produce DSBs as a consequence of cytokinesis [8–11]. Another condition that leads to MC concomitant with the appearance of anaphase bridges occurs when catenations between sister chromatids persist until anaphase. This happens when the catalytic action of topoisomerase II (Top2) is downregulated [12–14]. Top2 downregulation seems to pass undetected by cell cycle checkpoints in many cancer cell lines but not in normal differentiated cells [15–18]. Consequently, catalytic inhibition of Top2 offers a promising target to promote MC in cancer cells irrespective of the status of DNA damage checkpoints [19]. Of note, Top2 is often downregulated during acquisition of secondary resistance to chemotherapeutical regimes that comprise Top2 poisons, a major class of antitumor drugs [20, 21]. This observation implies that these resistant cancer cells should become even more hypersensitive to inhibition of Top2.

Top2 is essential in all organisms. In *Saccharomyces cerevisiae*, inactivation of Top2 by most thermosensitive (ts) alleles leads to MC in anaphase without previous checkpoint activation [13,22–26]. In recent years, *S. cerevisiae* has also been a model to study both cell death pathways and genomic instability footprints after environmental or genetic insults [27, 28]. Here, we have characterized the consequences for the offspring of the MC that follows inactivating Top2 through the *top2-5* ts allele (hereafter refer to as *top2* MC). We show that most of the *top2* MC progeny lose their ability to divide. Interestingly, these daughter cells do not die abruptly but undergo a slow decline in cell vitality over several hours. The patterns of cell death point towards an ACD, which was genetically corroborated with mutants for the main apoptotic pathway. We have also used heterozygous diploids to diagnose chromosome rearrangements in the surviving progeny, and we found genomic footprints that include uniparental disomy and terminal loss of heterozygosity in the longest chromosome arms. We conclude that (i) most *top2* daughter cells become senescent in the short-term while eventually dying by ACD; and (ii) the surviving offspring frequently carry genomic rearrangements expected from transiting through anaphase with intertwined sister chromatids.

## RESULTS

### Seventy five percent of the progeny of a *top2-5* mitotic catastrophe is inviable

We have recently reported that the *top2-5* thermosensitive mutant undergoes timely progression through the cell cycle until a MC occurs in late anaphase [25]. Importantly, *top2-5* gives a clear point-of-no-return in the MC phenotype because cytokinesis makes the *top2-5* anaphase bridges collapse irreversibly. In many ways, this MC is similar to other previously studied *top2* conditional alleles [13, 24], although *top2-5* provides a better synchrony for the MC since a larger percentage of cells quickly sever the anaphase bridge [25].

We performed single-cell videomicroscopy on agar plates through long-range objectives and found that mother and daughter cells struggled to rebud (the most obvious yeast signal for a new cell cycle) without Top2 (Fig. 1A) [25]. Whereas *TOP2* unbudded (G_1_/G_0_) cells were able to form microcolonies of around 10 cell bodies after 6 h at 37 °C, *top2-5* cells stopped dividing at either 2 (∼65%) or 3 (∼20%) cell bodies (Fig. 1A). We hereafter refer to cell bodies rather than cells or buds since it is difficult to conclude whether a 3 cell-body is part of a single multi-budded cell, a budded mother with a daughter, or a mother with two daughters. This 2-3 cell-body pattern was an end-point phenotype upon continuous Top2 inactivation, since we observed the same proportions after 24 h at 37 °C (Fig. 1B). Next, we investigated whether reactivation of Top2 by shifting the temperature down to 25 °C would allow any of these bodies to form a viable population. In order to have an overall picture of cell viability, we first determined clonogenic survival after different incubation periods at 37 °C. Because of the complexity of the budding patterns after the MC, we chose a solid medium-based clonogenic assay that allows to determine if at least one of the cell bodies was still viable by the time of the temperature shift, no matter how many cells are present in the progeny (Fig. 1C). We found that *top2-5* had a gradual loss of viability (50% survival after ∼ 4 h), and less than 5% clonogenic survival was obtained after 24 h at 37 °C (Fig. 1D); the *TOP2* isogenic strain retained the expected 100% clonogenic survival in this assay (Fig. S1A).

**Figure 1.**
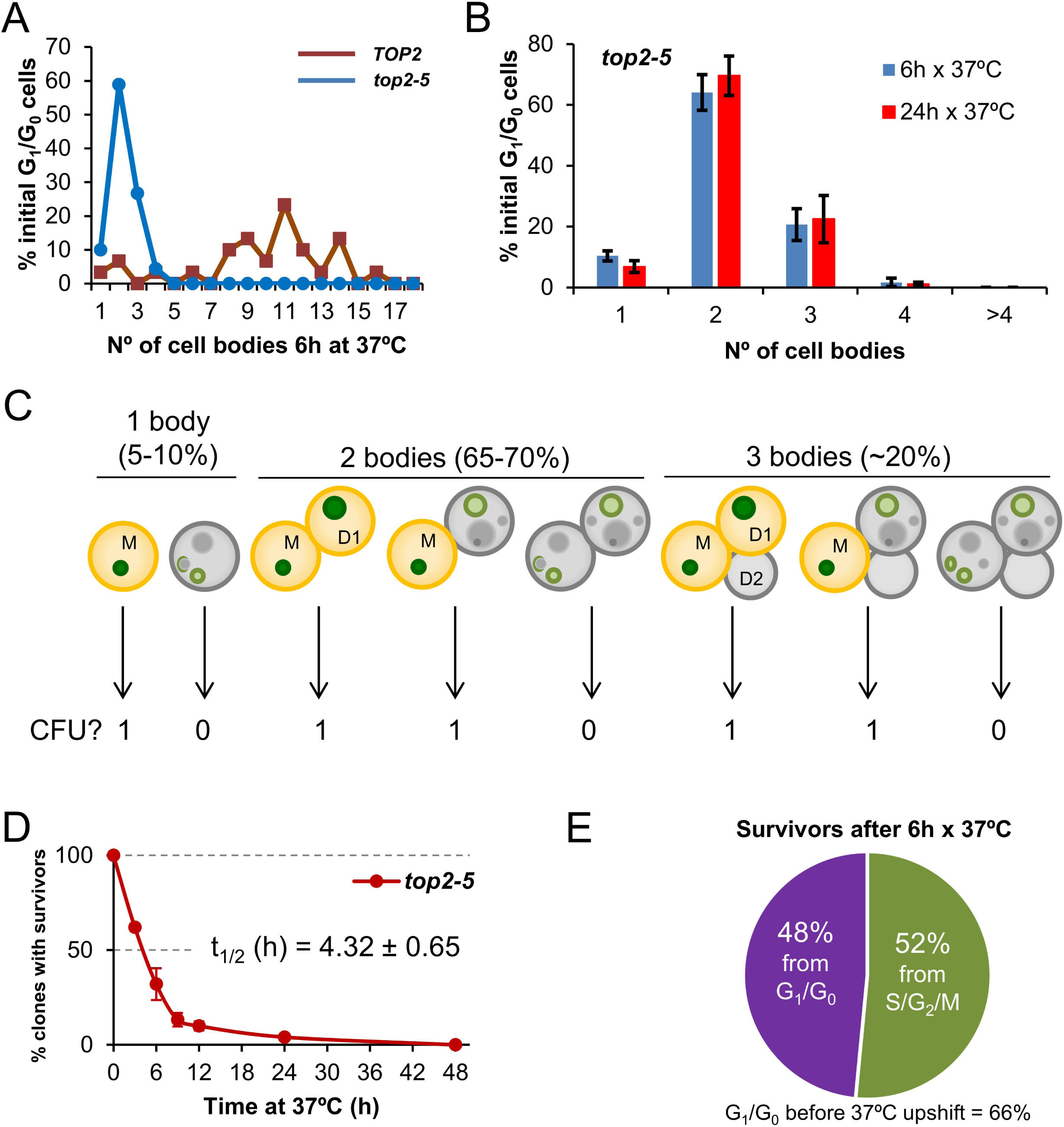
Most progeny coming from a Top2-mediated mitotic catastrophe is inviable. (**A**) Haploid *TOP2* (WT) or *top2-5* cells were grown at 25 °C and spread on YPD agar plates. Unbudded cells (G_1_/G_0_) were identified and photographed again after 6 h at 37 °C. Number of cell bodies (buds) coming from these G_1_/G_0_ cells were then counted and plotted as indicated. (**B**) The same analysis as in panel A but including data coming from independent experiments as well as after 24 h incubation at 37 °C (mean ± s.e.m., n=3). (**C**) The principle of the solid medium-based clonogenic assay. Unlike the liquid medium-based clonogenic assay, cells are spread on the Petri dish before the condition that challenges survivability is transiently triggered (Top2 inactivation in our study). In the solid medium-based assay, the colony forming unit (CFU) reading after the challenge is binary, irrespective of how far cells keep on dividing during the challenge: “0” if all clonal cells are inviable (grey); “1” if at least one cell from the clone stays viable (yellowish orange). (**D**) Time course of clonogenic survivability. Asynchronous *top2-5* cultures growing at 25 °C were spread onto several YPD plates. The plates were incubated at 37 °C for different periods before transferring them 25 °C. Four days after the initial plating, visible colonies (macrocolonies) were counted and normalized to a control plate which was never incubated at 37 °C (0h). (**E**) Analysis of the origin of macrocolonies after the 6 h x 37 °C regime as determined after microscanning plates at the time of seeding (N=33 macrocolonies; 2:1 unbudded:budded ratio at seeding).

Because in these clonogenic assays there is a mixture of budded (S/G_2_/M) and unbudded (G_1_/G_0_) cells at the time of plate seeding, we repeated the clonogenic survival after 6 h at 37° C, but photomicrographing the plate surface at different time points. Through this analysis, we determined that at time 0 h the unbudded:budded ratio was 2:1; however, only half of the surviving macrocolonies came from unbudded cells (Fig. 1E). This result implies that the chance to become a macrocolony is doubled if the original cell was budded at the time of the temperature upshift. The most likely explanation for this bias resides in the fact that the *top2* MC is expected to be milder if cells are closer to anaphase onset when Top2 is inactivated. Indeed, budded cells appeared to better complete the first two cell cycles at the restrictive conditions (Fig. S2). A calculation based on the cell proportions at 0h (66% G_1_/G_0_; 33% S/G_2_/M), overall macrocolony formation from the 37 °C for 6h regime (∼25%), and the origin of those macrocolonies (∼50% from G_1_/G_0_; ∼50% from S/G_2_/M) led us to conclude that around ∼20% of the original G_1_/G_0_ cells gave rise to survivors after 6 h at 37 °C [0.25 × 0.5 / 0.66 = 0.19]. This proportion increases to ∼40% for the S/G_2_/M cells at the time of the temperature upshift [0.25 × 0.5 / 0.33 = 0.38].

### Most of the inviable progeny of a *top2-5* mitotic catastrophe immediately stops dividing

Next, we tried to correlate the long-term clonogenic survival of the G_1_/G_0_ cells with their ability to form microcolonies. We reasoned that it is possible that many MCs could render viable progeny in the short-term (microcolony) but could not raise a visible colony later (macrocolony). This difference could reflect a gradual loss of viability in the progeny as a consequence of genomic imbalances acquired after the MC. To get further insights into this possibility, we took pictures of the cells on the surface of Petri dishes at the time of seeding (0h), right after the 37 °C incubations (6h or 24h), and 16 h or 24 h after the plates were shifted back to 25 °C (Fig. 2A). As expected from above, most of the original G_1_/G_0_ (unbudded) cells did not go beyond the 2-3 bodies stage during the 37 °C incubations (Fig. 2A, B; inner circles in the sunburst plots). Restoring permissive conditions for Top2 activity allowed very few of the cell progeny to divide again (Fig. 2A, B; outer circles in the sunburst plots). This finding was true not only during the long (24 h) incubation at 37 °C, but also for the 6 h incubation. Indeed, only ∼15% of the 2-3 cell bodies observed after 6 h at 37 °C were able to re-bud again once or more after shifting them back to 25 °C. Incidentally, a low but significant proportion of G_1_/G_0_ cells did not bud during the 37 °C regimes. However, even in these non-MC cases, cells did not divide after the Top2 reactivation, suggesting that this G_1_/G_0_ subpopulation was already incapable of cell division following growth at 25 °C. Longer incubations at 25 °C after the MC resulted in microcolonies of >50 cells that eventually developed into macrocolonies. With our cell density settings, microcolonies of more than 20-30 cells hindered us from raising conclusions about the fate of adjacent cells. However, the position of the center in these microcolonies suggests that most, if not all, of the G_1_/G_0_ cells that ended up as macrocolonies had re-budded again within the first 24 h that followed the temperature downshift.

**Figure 2.**
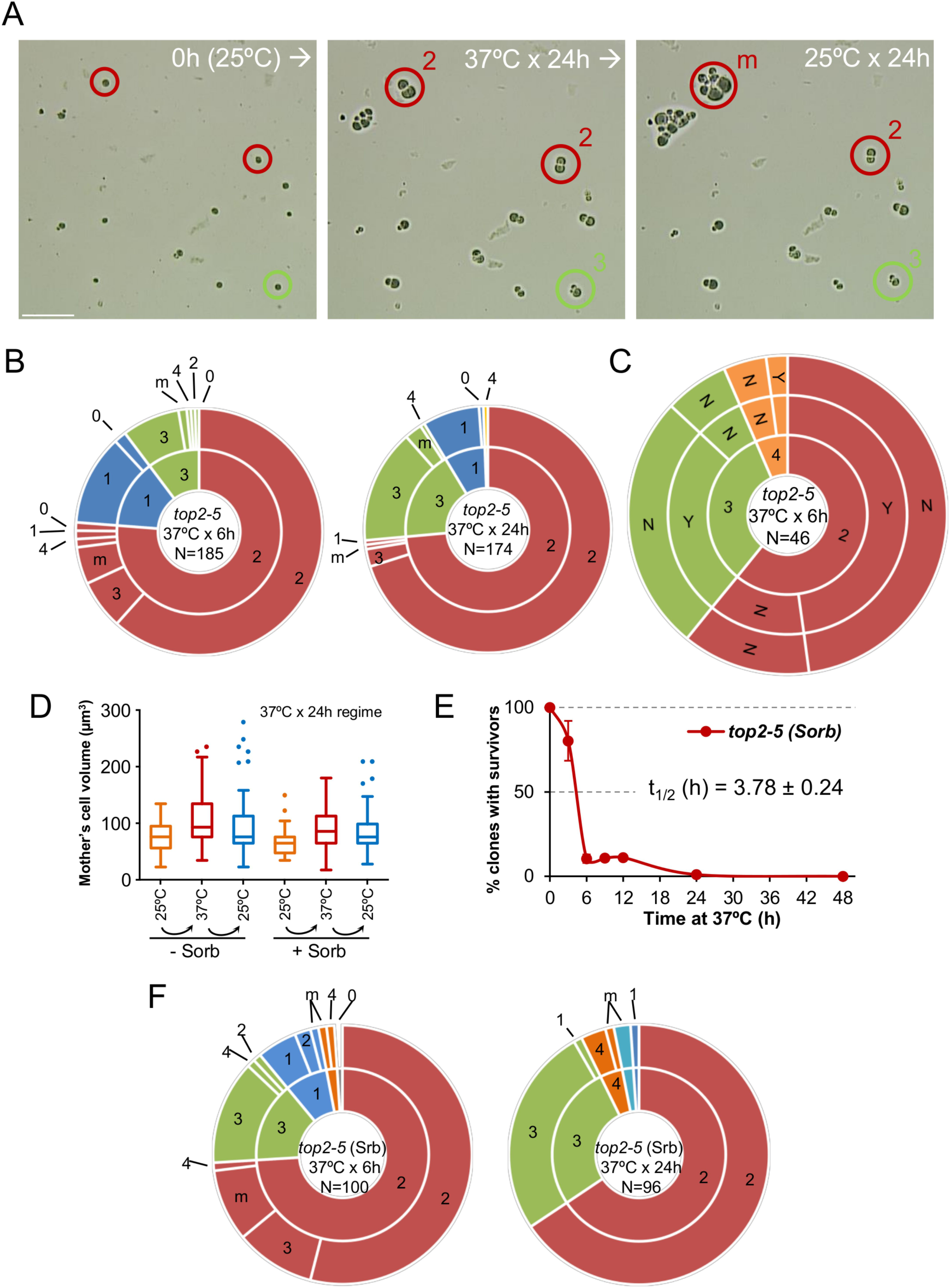
Most daughter cells coming from a Top2-mediated mitotic catastrophe are unable to divide again. (**A**) Haploid *top2-5* cells were spread at high cell density on two Petri dishes. At the time of seeding, 0h (25 °C), several fields were photomicrographed before incubating the plates at 37 °C during either 6 h or 24 h. After the 37 °C incubations, the same fields were localized, photomicrographed again, and further incubated 16-24 h at 25 °C. An example of a microscope field of a 37 °C x 24 h experiment. Three representative unbudded cells at 0h (25 °C) are highlighted. In red, two cells that budded just once during the 37 °C x 24 h incubation (“2” cell bodies); one of them able to re-bud again a few times after the 25 °C downshift (“m”) and the second one that remained stuck as “2”. In green, a cell that reached “3” bodies at 37° C and remained so after the final 25 °C x 24 h incubation. Scale bar corresponds to 50 μm. (**B**) Analysis of how far the *top2-5* MC progeny can go based on the microcolony approach shown in panel A. Only unbudded (G_1_/G_0_) cells at 0h (25 °C) were considered. The inner circle in the sunburst chart depicts proportions of cell bodies after the 37 °C incubation. The outer circle depicts the situation after the final 25 °C incubations (see supplemental information for a detailed description). On the left are results from a 37 °C x 6 h regime; on the right are results from a 37 °C x 24 h regime. Numbers point to the number of cell bodies; “m” means microcolonies of 5 or more bodies. (**C**) Capability of the *top2-5* progeny to split apart and relationship with overall survivability. Unbudded cells were micromanipulated and arranged at defined plate positions before incubating them for 6 h at 37 °C. Then, those cells able to re-bud at least once were subjected to an attempt to physically separate the cell bodies. The inner circle in the sunburst depicts the number of cell bodies after the 37 °C incubation. The middle circle depicts the result of the separation attempt (“Y” or “N”, successful or unsuccessful, respectively). The outer circle indicates if any of the bodies was able to raise a macrocolony (Yes or No) after 4 d incubation at 25 °C. (**D**) Progression of the size (volume) of the original G_1_/G_0_ cells (mother) after the *top2-5* mitotic catastrophe with and without the osmotic stabilizer Sorbitol (Sorb, 1.2 M). (**E**) Time course of clonogenic survivability in the presence of 1.2 M Sorbitol. The experiment was conducted as in Figure 1D. (**F**) Sunbursts of microcolony analyses in the presence of 1.2 M Sorbitol (Srb) at the 6 h and 24 h x 37 °C regimes. Interpretation as in panel B. In sunburst charts, N indicates number of original unbudded cells which were followed; blue sectors depict G_1_/G_0_ cells that remained unbudded during the 37 °C incubations; red sectors, cells that budded once at 37 °C; green sectors, cells that reached 3 bodies at 37 °C; orange sectors, cells that reached 4 bodies at 37 °C; cyan sectors, cells that reached 5 or more bodies at 37 °C.

From previous analysis by videomicroscopy, we know that most *top2-5* doublets (2 bodies) and all triplets (3 bodies) have passed anaphase and thus completed a MC after 6 h at 37 °C [25]. Nevertheless, we decided to complete the analysis of the immediate progeny by testing whether the observed cell bodies have accomplished cell separation. Our reasoning was that separation by micromanipulation would demonstrate that cell bodies have become individual daughter cells. Indeed, we could separate with the needle more than half of the doublets and triplets (Fig. 2C, middle concentric circle in the sunburst chart). We next tried to correlate the ability of all these cells to form macrocolonies but found that they were largely unviable. The percentage of macrocolonies (∼2%) was much lower than expected, indicating that that micromanipulation killed cells that otherwise would have retained viability.

We noticed from the microcolony experiments performed above that cells swelled after prolonged incubations at 37 °C (Fig. 2A). After 24 h at 37 °C, the volume occupied by the original mother cell doubled (Fig. 2D). Downshift of the temperature to 25 °C only modestly deflated these cells. In addition, a low proportion of cell bodies underwent lysis (“0”; 2 → 1 and 3 → 2 categories in Fig. 2B sunburst charts). These findings, together with the hypersensitivity to micromanipulation, led us to consider osmotic stress (due to the continuous growth in size of the mother cell) as a possible cause underlying the long-term inviability and/or short-term inability to divide of the *top2* progeny. However, addition of 1.2 M sorbitol, an osmotic stabilizer, neither prevented cells from swelling (Fig. 2D) nor improved long-term viability (Fig. 2E; S1B) or short-term division capability (Fig. 2F; Table S1).

### Cell death after prolonged absence of Top2 activity occurs slowly, asynchronously, asymmetrically and is independent of the Yca1 metacaspase

Next, we analysed cell bodies in these microcolonies more closely, seeking other morphological patterns of cell disease aside from swelling; for example, lysis, darkening and loss of the rounded shape (Fig. 3A). Most of these morphological features have been previously related to different forms of cell death. Because the *top2-5* strain also carries a GFP-labeled H2A histone, we also monitored nuclear morphology and chromatin integrity (i.e., GFP intensity). Cells still looked fully healthy after 6 h at 37 °C (no difference with 0 h), despite the great loss in the ability of the progeny to divide. Only after prolonged 37 °C incubations did the cells start to look clearly sick. Still, more than 50% of cell bodies harboured a nuclear GFP signal even 24 h following the 37 °C upshift. This GFP signal co-existed in cell progenies that showed unhealthy patterns in the bright field such as swelled, darkened, or non-rounded cell bodies.

**Figure 3.**
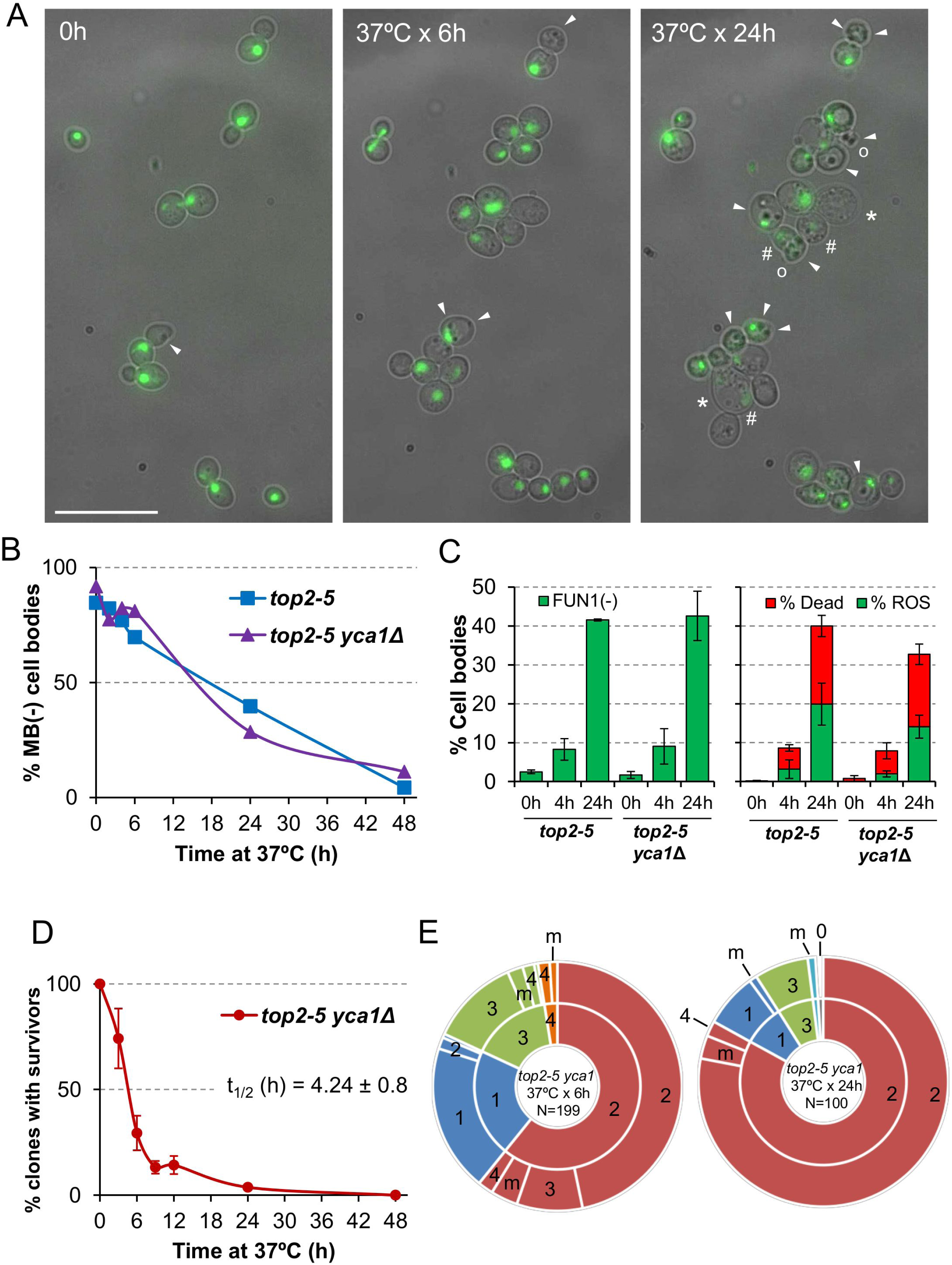
Cell vitality remain high for several hours after the *top2* mitotic catastrophe and is not modulated by Yca1. (**A**) Morphological patterns of cell and nuclear sickness after the *top2* MC. Haploid *top2-5 HTA2-GFP* cells were seeded onto agarose patches and the same fields visualized under the fluorescence microscope at 0 h, 6 h and 24 h after the 37 °C temperature upshift. White filled triangles point to darkened inclusion bodies, asterisks (*) swelled cells, open circles (○) cells that has lost their rounded shape, and hash (#) points to cells that have largely lost the H2A-GFP signal. BF, bright field. Scale bar corresponds to 20 μm. (**B**) Time course of cell vitality decline as reporter by methylene blue (MB) negative staining. Asynchronous cultures of the *top2-5* and *top2-5 yca1*Δ strains were grown at 25 °C before shifting the temperature to 37 °C. At the indicated time points (0, 2, 4, 6, 24 & 48 h), samples were taken and stained with the vital dye MB. (**C**) Cell vitality decline as reported by metabolic competence, intrinsic ROS generation, and loss of plasma membrane impermeability. Cells were treated as in B and stained at the indicated time points with the vital dye FUN1, the death marker propidium iodide (PI), and/or the ROS reporter DCFH-DA (mean ± s.e.m., n=3). (**D**) Clonogenic survival profile of *top2-5 yca1*Δ as determined on the low-density plates (mean ± s.e.m., n=3). The experimental procedure is described in Figure 1D. (**E**) Ability to re-bud of the *top2-5 yca1*Δ MC progeny as determined on the high-density plates. The experimental procedure is described in Figure 2.

In order to better study cell death after *top2* MC, we employed other means that required experiments to be performed in liquid media instead of Petri dishes. We first quantified the rate of cell death and metabolic decline by staining with the vital dye methylene blue (MB) (Fig. 3B). This dye stains dead cells blue, although it can also stain cells that are alive but metabolically attenuated [29]. A time course after the 37 °C upshift showed that there was not a major increase in MB positive cell bodies in the first 4 h (the equivalent in liquid cultures to 6 h on solid medium; [25]). In general, the increase in MB+ bodies was linear, but even 24 h after the 37 °C upshift ∼40% of cell bodies were not stained by MB. These staining experiments uncovered two other properties of the *top2* MC: i) it was common that only one cell body was MB+ in doublets and triplets (Fig. S3A); and ii) both unbudded and budded cells were equally stained (Fig. S3B). Regarding the first observation, the result suggests that loss of vitality is not coordinated between mother and daughter(s) cells. This asymmetry also confirms that many doublets must have completed cytokinesis despite remaining together. As for the second observation, the staining pattern suggests that loss of vitality occurs in an asynchronous fashion in terms of any preference for a cell cycle stage.

Because MB does not distinguish whether cell bodies are dead or simply metabolically stressed, we next sought other more informative vital stains. First, we employed the fluorescent vitality probe FUN1©. This probe stains metabolically active live cells with red vacuolar aggregates [30]. FUN1© is considered more informative and reliable than MB. With this probe we confirmed that only ∼10% of cell bodies have lost vitality after just 4 h at 37 °C (left chart of Fig. 3C; Fig. S4). It was also surprising that vitality decline still affected no more than 40% of all cell bodies after 24 h at 37 °C.

We also used Propidium Iodide (PI) to monitor cell viability. Loss of plasma membrane impermeability is considered a *bona fide* marker of cell death [27]. PI is only able to fluorescently stain cells that have lost such impermeability. Anticipating some sort of RCD after the *top2* MC, we decided to accompany PI staining in red with reporters for either reactive oxygen species (ROS) or externalization of phosphatidylserine (PS) at the plasma membrane; both in green (DCFH-DA and annexinV-FITC, respectively). Intrinsic ROS production has been observed during RCD in all eukaryotes, including yeast, and is considered one of most reliable RCD markers [31]. After overnight growth at 25 °C (0h), the *top2-5* strain had neither dead cells nor cells with ROS (right part of Fig. 3C, Fig. S5). Four hours after the 37 °C temperature shift, there was only a slight increase in dead cell bodies (∼6%) and almost no signs of ROS in the rest (∼3%). Only after 24 h of incubating the cells at 37 °C, we found a significant proportion of both ROS (∼20%) and dead cells (∼20%). It is noteworthy that the percentage of PI+ cells was still relatively low after this 24 h incubation, and 60% of all cell bodies were still resistant to PI and free of ROS. Similarly, PS externalization is considered a conserved *bona fide* marker of RCD [32]. Double staining with PI and annexin V-FITC in cells where the cell wall has been digested (a necessary step for annexin V to reach externalized PS) showed less than 10% of cell bodies with the staining expected for early apoptosis, even 24 h after the temperature upshift (Fig. S6; annexin V positive plus PI negative). It is noteworthy that there were more dead cell bodies (PI positive) in this assay than in the ROS/PI assay (∼60% vs. ∼20%). This observation is likely a consequence of digesting the cell wall, since the plasma membrane of swollen cells at 24 h might collapse during the treatment.

We next examined if the observed pattern of death after the *top2* MC was altered by a mutation in caspase. We reasoned that this approach might shed more light on the RCD/ACD nature of the observed death, considering the low presence of intrinsic ROS and the technical caveats of the annexin V assay. Hence, we deleted the only caspase-like gene in yeast, *YCA1/MCA1*. Yca1 is required for RCD in yeast in response to several environmental stresses [33–35]. In addition to cell death, we checked whether Yca1 modulated the other behaviours seen in the *top2* MC progeny (for example, the inability to divide and the slow decline in cell vitality). The conclusions derived from comparing *top2-5 YCA1* and *top2-5 yca1*Δ were that Yca1/Mca1: (i) had no influence in the percentage or rate of cells that end up dying (Fig. 3B, C); (ii) neither accelerated nor slowed down the vitality decline (Fig. 3B, C); (iii) did not modify the profile of clonogenic survival after the *top2* MC (Fig. 3D; S1C); and (iv) its absence did not improve the ability of the immediate cell progeny to divide (Fig. 3E; Table S1).

The overall conclusion from these experiments is that the immediate progeny of the *top2* MC enter a senescent-like state as they retain vitality but loses their ability to re-bud. Senescence is only a transient state that lasts several hours or days, until cells eventually die. The pattern of cell death (morphologically diverse, slow, asynchronous and asymmetrical), together with the lack of effect of Yca1, suggest that loss of Top2 leads to ACD.

### Chromosome ploidy modulates the ability of the progeny to divide

All the experiments described above were carried out in haploid yeast cells. In haploids, MC associated with the presence of partly unresolved sister chromatids, as in the *top2-5* mutant, is expected to be highly deleterious. Severing of these anaphase bridges result in daughter cells that may lack several chromosome arms [11]. Based on this consideration, one might expect that diploid cells would be more resistant to the consequences of *top2* MC than haploid cells. Thus, we studied the *top2* MC in an isogenic homozygous *top2-5/top2-5* diploid (2N) strain. Unlike its haploid counterparts, diploid cells were more often able to re-bud at least once. In fact, ∼50% of all 2-3 cell bodies originated from just after 6h at 37 °C were able to do so after the 25 °C downshift (Fig. 4A; Table S1). This percentage dropped considerably if the progeny was incubated 24 h at 37 °C. Strikingly, however, the increase in the ability to divide again after the MC did not yield better clonogenic survival (Fig. 4B; S1D), indicating that most of this viable progeny was competent to re-bud only in the first generations. We also performed a microdissection analysis of the diploid *top2* progeny. Unlike haploid *top2-5* cells, which was rather sensitive to micromanipulation, 13% of the diploid progeny raised a macrocolony after the separation attempt (Fig. 4C). Altogether, we conclude that a *top2* MC in diploids results in better short-term ability to divide.

**Figure 4.**
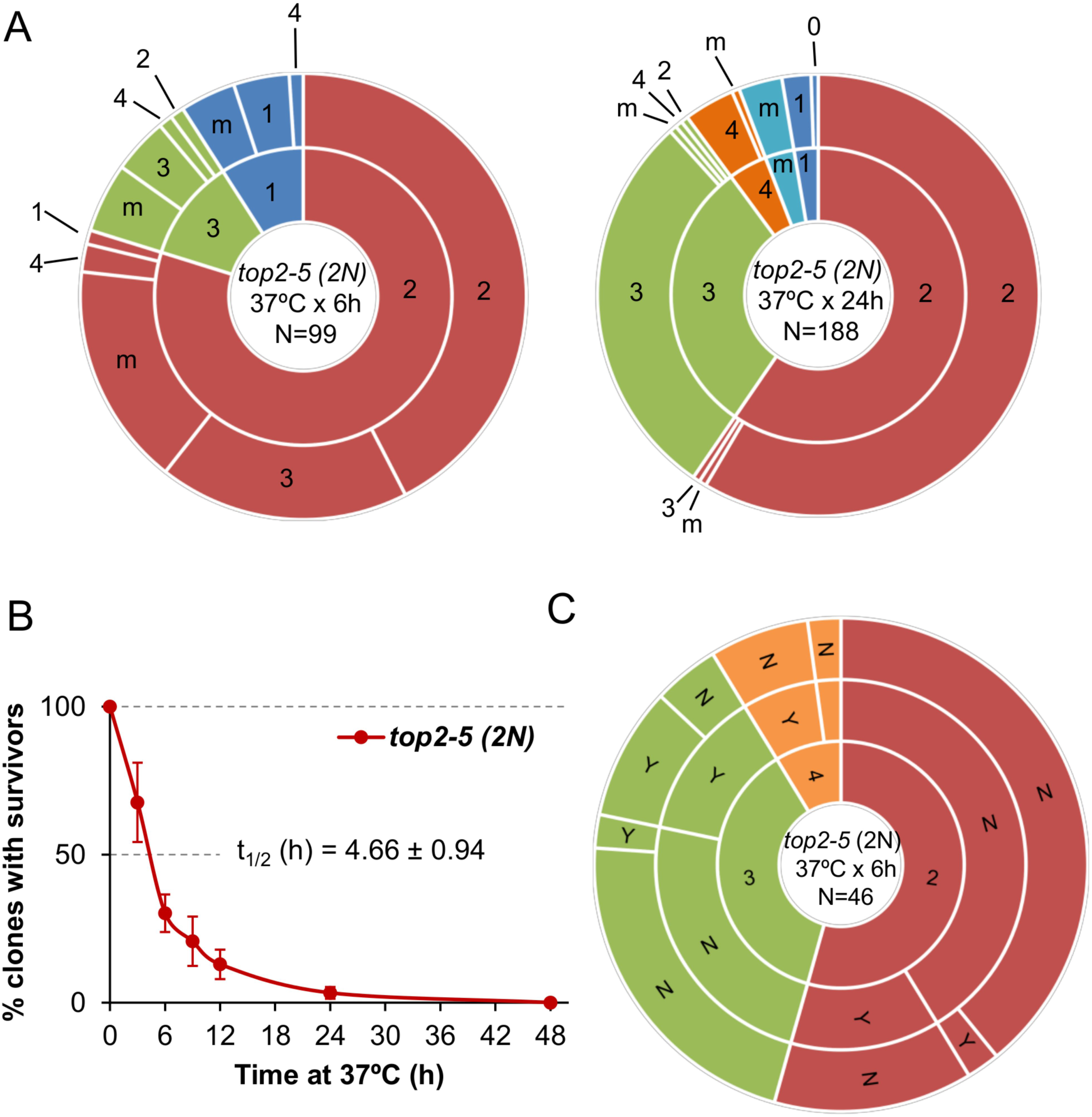
Mitotic catastrophe in *top2-5* diploids leads to progeny with a greater capacity for cell division than observed in the haploid. Isogenic homozygous *top2-5* diploid cells were grown and spread at either low or high cell density on Petri dishes. In addition, G_1_/G_0_ cells were micromanipulated, arrayed and treated as described in Figure 2C. (**A**) Ability to re-bud after transient (6 h or 24 h) incubations at 37 °C of the high-density plates. The experimental procedure is described in Figure 2. (**B**) Clonogenic survival profile as determined on the low-density plates (mean ± s.e.m., n=3). The experimental procedure is described in Figure 1D. (**C**) Capability of the progeny to split apart and relationship with overall survivability. The experimental procedure is described in Figure 2C.

### Genome instability footprints in the surviving progeny from the *top2*-mediated mitotic catastrophe

Above, we have just shown that ∼25% of the *top2-5* diploid cells still gave rise a macrocolony after the 6h at 37°C regime. We next examined the genomes of these survivors in search for specific genomic footprints of the *top2* MC. To accomplish this goal, instead of using the homozygous isogenic diploid employed in the previous chapter, we generated a highly heterozygous hybrid *top2-5* diploid. The hybrid diploid was generated by crosses of two sequence-diverged *top2-5* haploids, derivatives of W303-1A and YJM789, which are heterozygous for more than 55,000 single-nucleotide polymorphisms (SNPs) distributed throughout the yeast genome (the yeast genome is 15 Mb). These heterozygous SNPs allow the analysis of various types of genomic alterations by using SNP-specific microarrays [36, 37]. The hybrid diploid is also engineered to select various types of chromosome alterations on chromosome V (Fig. 5A). It is homozygous for the *ade2-1* mutation on chromosome XV (an ochre-suppressible allele) and heterozygous for the *SUP4-o* suppressor gene in chromosome V [38]. Strains with the *ade2-1* mutation form red colonies in YPD in the absence of the *SUP4-o* suppressor. Diploid strains with one or two copies of *SUP4-o* form pink and white colonies, respectively [39]. Thus, loss of the *SUP4-o* gene by either mitotic recombination or chromosome loss results in a red colony instead of the pink colonies characteristic of the original strain. Colony colour changes are also expected if the copy number of the *ade2-1* allele varies by loss or duplication of chromosome XV. In addition, aneuploidy for other chromosomes sometimes alters the color of the colony.

**Figure 5.**
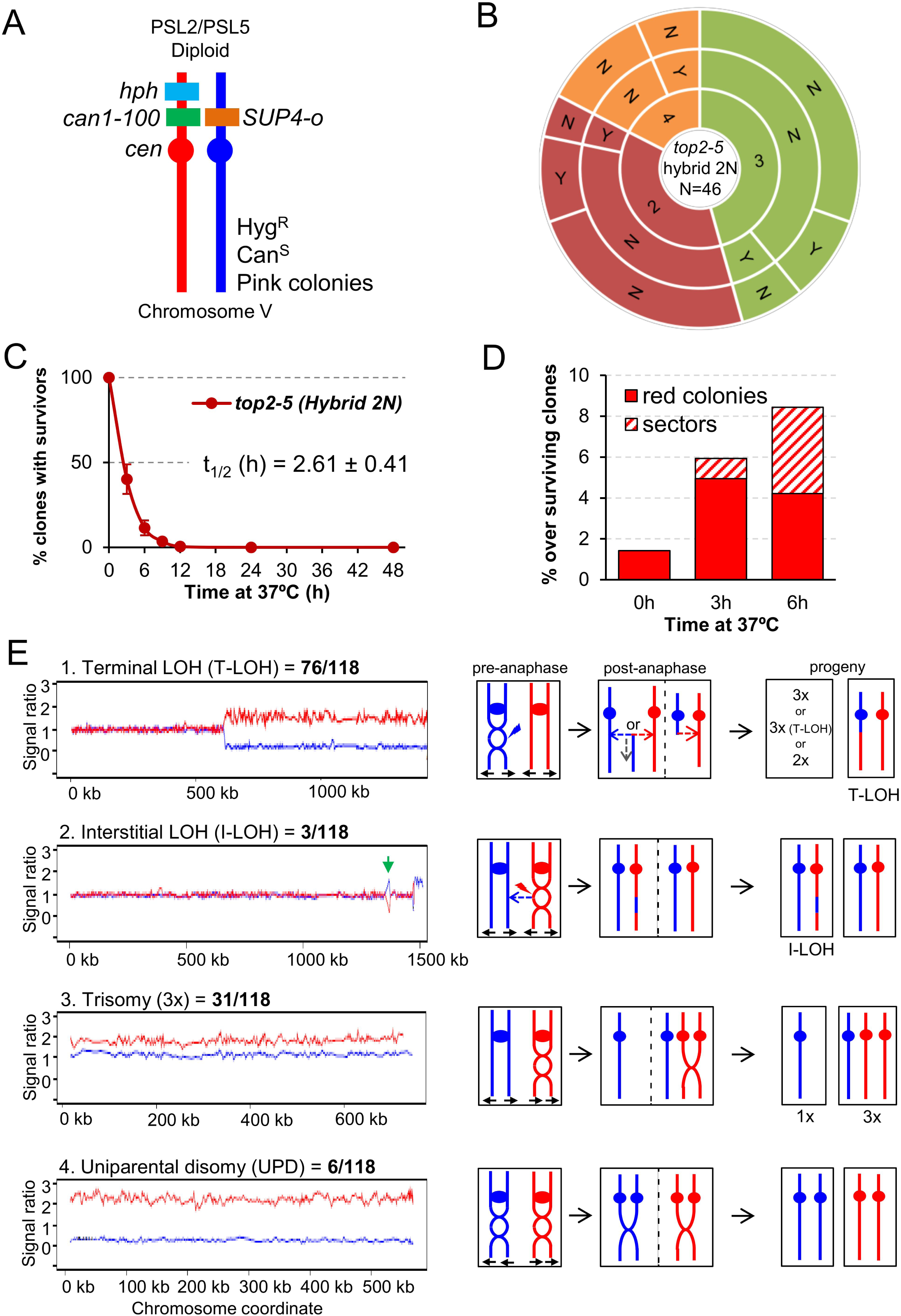
Mitotic catastrophe in *top2-5* heterozygous diploids leads to survivors with specific genetic instability footprints. (**A**) Schematic of the engineered chromosome V (cV) from the hybrid highly heterozygous (∼55,000 SNPs) diploids used in this study. As explained in the text, the genetic modifications applied in cV allowed for selection of chromosome rearrangements. (**B**) G_1_/G_0_ cells from the hybrid highly heterozygous *top2-5* diploid FM1873 strain were micromanipulated, arrayed and treated as described in Figure 2C. The capability of the immediate progeny to split apart and its relationship with overall survivability is shown in the sunburst chart. (**C**) Clonogenic survival profile of FM1873 as determined on low-density plates (mean ± s.e.m., n=3). The experimental procedure is described in Figure 1D. (**D**) Percentage of red or sectored (either white:red or pink:red) colonies in the surviving clones. Both outcomes often reflect genetic alterations on cV as described in the text. (**E**) Results of SNP microarray analysis of colonies derived from FM1873 or MD684. Microarray patterns showing specific chromosome rearrangements are shown on the left side, and diagrams of the putative events producing these patterns are shown on the right side. The number of specific events out of 118 total events is indicated. For the microarray patterns, hybridization to SNPs specific to homologs derived from W303-1A are shown in red, and hybridizations to SNPs specific to YJM789 are shown in blue. The X-axis shows SGD coordinates for the chromosome, and the Y-axis shows the ratio of hybridization normalized to a heterozygous diploid strain. The representative examples correspond to (1) a T-LOH event on chromosome IV (MD684.1.1 (E1) in Table 1); (2) a I-LOH event (marked with a green arrow) plus T-LOH event on chromosome IV (FM1873-15c (C2) in Table 1); (3) a Trisomy for chromosome XIV (MD684.1.1 (E1) in Table 1); and (4) a UPD for chromosome V (This isolate has two copies of the W303-1A-derived and no copies of the YJM789-derived chromosome).

**Table 1.** Genomic changes in single-colony isolates of FM1873 and MD684.

| **Strain name<sup>1</sup>** | **Genomic alterations<sup>2</sup>** |
| --- | --- |
| FM1873-01 (E1) | T-LOH (IV/966), Tri (V), Partial UPD (VII), Tri (X) |
| FM1873-04 (E1) | T-LOH (IV/471), UPD (VIII), 3 T-LOH (XIII/450/777/864) |
| FM1873-11 (E1) | UPD (IV), Tri (VIII), Mon (IX), Tri (X), Partial Tri (XV), UPD (XVI) |
| FM1873-1 (E2) | UPD (IV), Tri (XV) |
| FM1873-2 (E2) | T-LOH (IV/790) |
| FM1873-3 (E2) | T-LOH (IV/790) |
| FM1873-4 (E2) | T-LOH (IV/671), Tri (V), Tri (VII), T-LOH (XV/1002) |
| FM1873-5 (E2) | T-LOH (IV/755), T-LOH (XIII/450) |
| FM1873-11X (E2) | T-LOH (IV/958), Tri (VII) |
| FM1873-12 (E2) | Tri (VIII) |
| FM1873-13 (E2) | T-LOH (IV/668), Tri (V), T-LOH (XV/1002) |
| FM1873-14 (E2) | T-LOH (IV/1018) |
| FM1873-15 (E2) | T-LOH (IV/483) |
| FM1873-C1 (C1) | No additional alterations |
| FM1873-C2 (C1) | No additional alterations |
| FM1873-C3 (C1) | 2 T-LOH (IV/485/820) |
| FM1873-C4 (C1) | T-LOH (IV/470), T-LOH (XIII/856), T-LOH (XIV/216) |
| FM1873-1c (C2) | 2 T-LOH (IV/456/1440); T-LOH (VII/385), T-LOH (XIV/400) |
| FM1873-2c (C2) | 2 T-LOH (IV/957/1440) |
| FM1873-3c (C2) | T-LOH (IV/935), Tri (V) |
| FM1873-4c (C2) | T-LOH (IV/858) |
| FM1873-5c (C2) | Tri and T-LOH (IV/1270), Tri (V), Partial Tri (XIII) |
| FM1873-11c (C2) | T-LOH (IV/1017), Tri (VIII), Tri (XV) |
| FM1873-12c (C2) | T-LOH (IV/1004), Tri (X), Tri (XV) |
| FM1873-13c (C2) | T-LOH (IV/1017), Tri (VIII) |
| FM1873-14c (C2) | 2 T-LOH (IV/889/1221) |
| FM1873-15c (C2) | I-LOH (IV/1354-1357), T-LOH (IV/1474), Tri (V), UPD (VIII) |
| MD684.1.1 (E1) | T-LOH (IV/596), T-LOH (XI/7.5), T-LOH (XII/430) |
| MD684.1.2 (E1) | T-LOH (IV/808), Tri (VIII) |
| MD684.1.3 (E1) | T-LOH (IV/1002), T-LOH (XII/165) |
| MD684.1.4 (E1) | T-LOH (IV/458), Tri (XV) |
| MD684.1.5 (E1) | T-LOH (IV/1065), Tri (VIII), 2 T-LOH (XII/253/446) |
| MD684.1.6 (E2) | Partial Tri (II) |
| MD684.1.7 (E2) | T-LOH (IV/1028), T-LOH (VII/29), T-LOH (XII/175) |
| MD684.1.8 (E2) | 3 T-LOH (IV/921/1050/1070), T-LOH (V/538), T-LOH (XII/450) |
| MD684.1.9 (E2) | Partial Mon (I), Tri and T-LOH (IV/486) |
| MD684.1.10 (E2) | 2 T-LOH (IV/661/963), Tri and T-LOH (XV/626) |
| MD684 C1.1 (C1) | I-LOH (IV/747-760), T-LOH (IV/840), T-LOH (XII/176) |
| MD684 C1.2 (C1) | T-LOH (IV/458), T-LOH (XII/447) |
| MD684 C1.3 (C1) | 2 T-LOH (IV/893/1472), Tri (V), T-LOH (XII/170) |
| MD684 C1.4 (C1) | Tri (I), T-LOH (IV/920) |
| MD684 C1.5 (C1) | T-LOH (IV/527), Tri (XII) |
| MD684 C1.6 (C2) | 2 T-LOH (IV/680/1050), T-LOH (VII/760), T-LOH (XII/175) |
| MD684 C1.7 (C2) | T-LOH (IV/1008), I-LOH (XII/215-228), T-LOH (XII/277), Partial Tri (XVI) |
| MD684 C1.8 (C2) | T-LOH (IV/842), T-LOH (XV/657) |
| MD684 C1.9 (C2) | T-LOH (IV/1028) |

<sup>1</sup> Parentheses after the strain name indicate whether the strain was experimental (E, incubated for six hours at 37 °C) or control (C, not incubated at 37 °C). E1 and C1 indicates that the 37 °C incubation was done on plates; in E2 and C2 experiments, the 37 °C incubations were done in liquid.
<sup>2</sup> Strains derived from FM1873 (both experimental (incubated at 37 °C for six hours) and control (not incubated at 37 °C) strains had three to four copies of chromosome XIV and a terminal LOH event on the right arm of chromosome XII (breakpoint at 236 kb). These alterations, therefore, are not listed in the FM1873 strains. The MD684 strain also had three to four copies of chromosome XIV in isolates, and this alteration is not shown in the table. Code: T-LOH (terminal LOH event), I-LOH (interstitial LOH event), Tri (trisomy), Mon (monosomy), UPD (uniparental disomy). Between brackets: altered chromosomes/breakpoints.

Microcolony and clonogenic experiments performed in the *top2-5* hybrid diploid FM1873 showed that this strain lost viability quicker than the isogenic homozygous diploid in the S288C background (Fig. 5B, C; S1E). There was a steady rise in red and/or red/white sectored colonies among the survivors during the 37 °C incubation, as expected if the *top2* MC increases genome instability (Fig. 5D). We used microarrays to examine genomic alterations in the control FM1873 isolate (no exposure to 37 °C) and in isolates exposed to 37 °C for 6 h, and then allowed to form colonies at 25 °C (Fig. 5E; see also supplemental information for interpretation on the genomic alterations picked up by SNP array). It is important to stress that the SNP microarrays allow analysis of genomic alterations throughout the genome in addition to those changes that occur on chromosome V [36, 37]. When the control FM1873 strain was examined before exposure to the restrictive temperature, surprisingly, we found that it was altered relative to an isogenic *TOP2*/*TOP2* hybrid [40]. More specifically, we realized that FM1873 carried a terminal loss of heterozygosity (T-LOH) on the right arm of chromosome XII (the longest chromosome arm in yeast). In addition, all isolates had three to four copies of chromosome XIV. We tried several times to recreate the hybrid *top2-5* diploid but were unable to isolate a derivative that had only two copies of XIV. Since this chromosome is the location of *top2-5*, it is likely that chromosome XIV trisomes and tetrasomes have a selective growth advantage at the permissive temperature over the diploids that have only two *top2-5* copies. We generated an isogenic derivative of FM1873 (MD684) that lacked the T-LOH event on XII, although it still had extra copies of XIV. For our subsequent genomic analyses, we studied both FM1873 and MD684.

The experimental strains were exposed to the restrictive temperature of 37 °C by incubating the cells for six hours either on plates or in liquid (details in Materials and Methods); these two protocols resulted in similar levels of instability. A total of 27 isolates were examined for FM1873 (13 experimental, 14 control), and 19 isolates of MD684 (10 experimental, 9 control). Somewhat surprisingly, the control single-colony isolates (cells not exposed to 37 °C) also had high rates of instability (Table 1; “C” samples), indicating that the Top2p encoded by *top2-5* does not have wild-type activity even at the permissive temperature. Indeed, a previous biochemical study reported that the Top2-5 activity at 25 °C is 33% of that of wild type Top2 [41]. Among all isolates examined, we found 76 T-LOH events, 3 interstitial LOH (I-LOH) events, 31 trisomies, 2 monosomies, and 6 uniparental disomies (UPDs) (Fig. 5E; Table 1). The average number of genetic changes per strain (including all the data of Table 1) was 3.3 alterations/isolate. Since the strains were grown approximately 40 generations before microarray analysis, the rate of alterations/cell division/isolate is about 8.3 x 10^-2^. In a previous microarray analysis of a wild-type diploid isogenic with FM1873 and MD684, we found a rate of alterations of about 2 x 10^-3^/division, a rate about 40-fold less than for the *top2* strains. The most common alteration in the *top2* strains was a T-LOH event (64% of the total events). These events likely reflect the repair of a DSB by either a crossover or a break-induced replication (BIR) event (Fig. 5E). The chromosomal distribution of the events is striking. The right arms of chromosome IV and XII (the two longest arms in the yeast genome) had over 80% of the terminal LOH events (60/76), although these arms represent less than 30% of the yeast genome. Other chromosomes with T-LOH events (the number of events shown in parentheses) are: XIII (5), XV (4), VII (3), XIV (2), VII (1), XI (1), and V (1). Chromosomes XIII, XV and XIV have the third, fourth, and fifth longest chromosome arms in the genome, respectively; all of the mapped events in these chromosomes are on the longest arms. Strikingly, the very large (about 1.2 Mb) ribosomal RNA gene cluster (rDNA) on the right arm of chromosome XII was not a preferred site for T-LOH events. Although the rDNA is about 60% of the right arm of XII, only 30% (3 of 11) of the LOH events had a breakpoint in or near the rRNA genes. We should point out that T-LOH events on XII could only be followed in the MD684 strains (FM1873 already had a cXIIr T-LOH); hence, our estimate of the frequency of T-LOH events on the right arm of XII is a minimal estimate.

In addition to LOH events, we observed 33 changes in chromosome number (31 trisomies and 2 monosomies) (Table 1). The frequencies of trisomies are not simply related to the size of the chromosomes. Chromosomes V, VIII, and XV were the most frequently-observed trisomes; chromosomes V and VIII are medium-sized chromosomes (both about 550 kb), whereas chromosome XV is large (1092 kb). Only one trisomy was observed involving one of the three smallest chromosomes (I, VI, and III). Only three of the trisomies involved the two largest chromosomes IV and XII.

We also observed six UPD events (Fig. 5E). In strains with these events, the homolog is present in two copies, but both copies are derived from one of the original parental homologs. There are two plausible pathways to generate UPD (Fig. S7): two non-disjunction events in different cell cycles or reciprocal UPD (RUPD) in which one pair of homologs segregates into one cell and the other pair segregates into the other cell. Although both pathways probably contribute to UPD in yeast, at least some of the events in wild-type strains are RUPD [42]. To determine whether RUPD events occurred frequently in the *top2* cells, we used a protocol in which both daughter cells produced as a result of RUPD or a reciprocal crossover (RCO) in chromosome V could be recovered in different sectors of a sectored colony (Fig. S8). The sectored colonies were derived from the same strains (FM1873 and MD684) used for our single-colony analysis. A crossover or RUPD can produce a red/white sectored colony. However, to select for such events, both FM1873 and MD684 contained a heterozygous *can1-100* marker located allelically to the *SUP4-o* insertion and a gene encoding hygromycin resistance (*hph*) located distal to the *can1-100* insertion (Fig. 5A). The *can1-100* mutation is a nonsense mutation and is suppressed by *SUP4-o*. Strains that lack the suppressor are resistant to the drug canavanine and those with the suppressor are sensitive. A crossover between *can1-100* and the centromere results in formation of a canavanine-resistant red/white sectored colony (Fig. S8A); a sectored colony with the same phenotype can also result from RUPD (Fig. S8B). RCO and RUPD events can be distinguished by microarray analysis (bottom panels of Fig. S8).

Following exposure of FM1873 and MD684 to 37 °C, we found 242 red/white Can^R^ sector colonies, 43 of which had the sectoring pattern for the *hph* marker suggestive of RCO or RUPD events. We found both RCO and RUPD events in two-thirds of these colonies (Table S2). The rate of RUPDs in cells of FM1873 and MD684 treated for 6 h at 37 °C was the same, 1.1 x 10^-5^/division; the rate of RUPD in cells of FM1873 that were not exposed to 37 °C was 1 x 10^-6^. The rate of RCO in FM1873 cells treated at 37 °C was 2.7 x 10^-6^; no RCO events were observed in MD684. In an isogenic wild-type strain, the rates of RUPD and RCO for chromosome V were 10^-7^ and 1.6 x 10^-6^, respectively [42]. These results support the conclusion that the *top2* mutation results in a substantially elevated rate of RUPD (about 100-fold) and has little effect on the rate of RCO.

## DISCUSSION

The formation of anaphase chromosome bridges during the cell division is one of the most dramatic sources of genetic instability. Such bridges are expected to cause a MC that would kill most of the progeny. Downregulation of Top2 is likely the most common way of generating large numbers of anaphase bridges. In addition, downregulation of Top2 has important clinical implications during acquisition of resistance against cancer therapy that comprises Top2 poisons (for example, etoposide and anthracyclines). In the present work, we have studied the consequences of inactivating Top2 in yeast through the *top2-5* thermosensitive allele. We show that the expected MC often leads to progeny unable to divide; however, cell death is not immediate but the result of a decline in cell vitality that takes hours to complete. We further propose that the irreversible genomic imbalances that occur during chromosome segregation in the absence of Top2 explain the short-term senescence observed in the immediate progeny. This hypothesis is strengthened by the observation that diploid survivors of a *top2* MC often carry genomic footprints expected from anaphase bridges such as UPDs and T-LOH. A step-by-step summary of the *top2* MC events is shown in Fig. 6.

**Figure 6.**
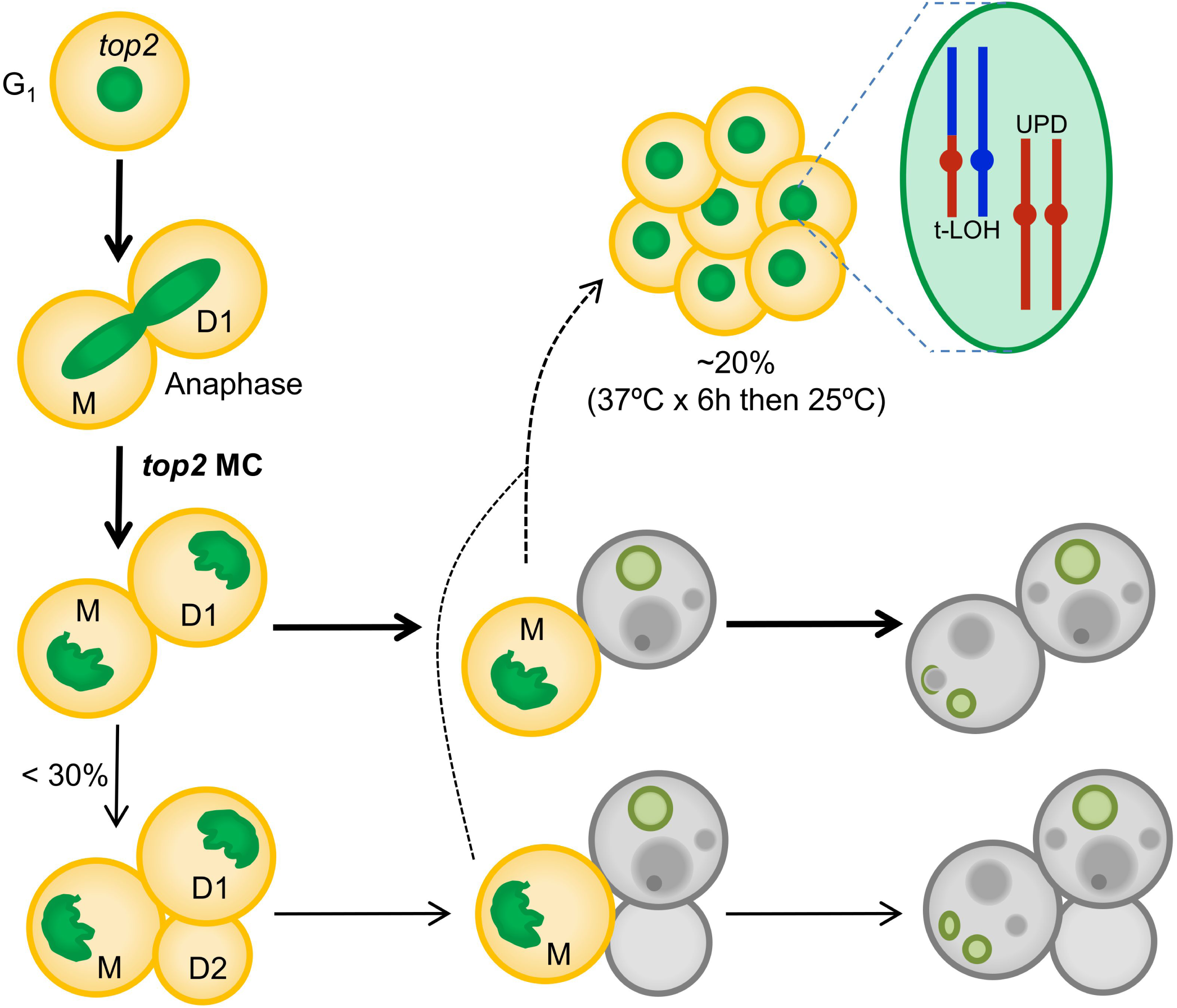
Summary of the *top2*-mediated mitotic catastrophe and the fate of the immediate progeny. After inactivation of Top2, cells cannot resolve sister chromatids in anaphase, leading to an anaphase bridge between the mother (M) and its daughter (D1). These bridges are quickly severed (at least in the *top2-5* mutant [25]). The immediate progeny coming from the *top2* mitotic catastrophes (MCs) is largely unable to enter a new cell cycle (do not re-bud) despite remaining metabolically active for many hours; hence, these cells become senescent. Only ∼25% of the original mothers re-bud once (D2) after the *top2* MC. The long-term fate of most daughter cells is death. They will eventually die through accidental cell death (ACD), as deduced from both the asynchrony and asymmetry of death events and the lack of regulation by the death modulator Yca1(Mca1). The inability to enter a new cell cycle is likely a consequence of both the massive DNA damage as a consequence of bridge severing, and the misdistribution of essential genetic material coded on the chromosome arms between the daughter cells. A small proportion of the progeny, especially those cell that underwent a milder *top2* MC (e.g., already in S/G_2_ at the time of Top2 inactivation) survives to yield a population of cells with characteristic footprints of genomic instability. Two of these footprints, terminal loss of heterozygosity (T-LOH) and uniparental disomy (UPD) are expected outcomes from anaphase bridges.

We would like to point out there are many different strategies to abolish Top2 activity, from depleting the enzyme through degron systems to using catalytically dead mutants. In between, there are a large collection of conditional thermosensitive alleles. Baxter and Diffley [24] showed that depletion of Top2 had a different phenotype than expression of a catalytically-dead enzyme. Depletion of Top2 did not lead to cell-cycle delays but did result in lethal DNA damage during cytokinesis. In contrast, a mutation in the Top2 catalytic domain resulted in arrest in G_2_/M as a consequence of DNA damage (nicks and gaps). The absence of a G_2_/M arrest is also a common pattern of thermosensitive alleles [13,22–26], with the remarkable exception of top2-B44 (F977L), which delays cells in G_2_/M through activation of the spindle assembly checkpoint [26]. In addition, we must consider post-G_2_/M delays since the NoCut checkpoint slows down cytokinesis under the presence of anaphase bridges [43]. This was shown using the *top2-4* allele, and this delay affects the synchrony of the MC. We previously examined the kinetics of both completion of cytokinesis and severing of anaphase bridges in *top2-4* and *top2-5* [25]. We found that the *top2-5* strain severed the bridge more efficiently and synchronously than *top2-4*, arguing that *top2-5* is a better allele to model *top2* MC; hence, we used this allele to address the *top2* MC. It would be interesting, though, to address the consequences of *top2* MC in other models, including degron-assisted Top2 depletion, partial inactivation (semipermissive conditions) and *top2-ts* alleles that delay either G_2_/M or cytokinesis.

### On the short-term cytological consequences of the *top2* mitotic catastrophe

In this study we have adopted the term mitotic catastrophe (MC) in its broadest cytological sense, referring to aberrant mitoses that are expected be deleterious for the progeny based on the degree of observed abnormalities. It is worth mentioning that other authors, especially those working with metazoans, restrict the MC term to death occurring in mitosis after a mitotic insult [44, 45]. With this restriction in mind, MC in metazoans is a sort of mitotic RCD. In our yeast experimental model with the *top2-5* allele, death before anaphase is insignificant since cells go through G_2_/M and anaphase as quickly as their *TOP2* counterparts [25]. In nocodazole-treated cells, which arrest cells in G_2_/M, 40% cell death was reported after 10 h in clonogenic assays [46]. This cell death was described to occur through an RCD apoptosis-like mechanism. By contrast, our clonogenic assays showed that a sudden drop in viability occurs between 3-6 h after Top2 inactivation (Fig 1C), about the time needed for cells to complete telophase and cytokinesis on solid medium [25]. From the microcolony experiments, we concluded that the ensuing *top2-5* progeny are largely impaired in entering a second cell cycle (Fig 1A, B and 2). This impairment can be partly alleviated in diploids, yet only in the short-term (Fig 4). This observation leads us to propose that gross genomic imbalances prevent the immediate progeny from cell cycle progression. Taking into account previous reports on the formation of anaphase bridges in *top2-ts* mutants [13, 25] and high levels of chromosome missegregation [23], it appears logical that many of the haploid progeny lose entire chromosomes containing essential genes. In addition, loss of essential genes may reflect breakage of chromosomes at the bridge followed by loss of the distal chromosome regions [11].

Another possibility is that daughter cells immediately die upon the *top2* MC through an RCD program. This would imply that either anaphase/telophase cells or their immediate progeny sense the MC and execute an RCD. Our results, however, argue against this possibility. Firstly, many cells stained negative for death (PI) and apoptotic (annexin V) markers, whereas they stained positive for metabolic activity (FUN1©), even after 24 h at 37 °C. Secondly, the decline in cell vitality occurred slowly, linearly (asynchronously) and asymmetrically (when comparing daughter cells that remained together after the MC). Thirdly, Yca1/Mca1, the main RCD player in *S. cerevisiae*, does not regulate the vitality declines. Even though there are other RCD proteins aside from Yca1/Mca1 [47, 48], we point out that that the pattern of cell death after the *top2* MC is better explained through ACD. Although we found intrinsic ROS production in a minor subset of the *top2* MC progeny, and this finding has been considered a marker of RCD [31, 49], we hypothesize that, in our case, ROS accumulation is a consequence of the steady decline of cell homeostasis. For instance, ROS might arise from loss of nuclear genes involved in eliminating ROS in metabolically active cells.

Comparing with previous studies, the events that lead to death after the *top2* MC resemble those observed after prolonged G_2_/M arrest in the *cdc13-1* mutant, which results in irreversible DNA damage at chromosome ends. In arrested *cdc13* cells, there are cell markers of RCD such as ROS production [50], although further biochemical assays and genetic manipulation suggest ACD rather than RCD [47]. In *cdc13-1*, because cells get blocked in G_2_/M, there are no genetic imbalances prior to cell death. It was proposed that cell lysis, resulting from cell growth without cell division, was the ultimate cause of death, a hypothesis confirmed because sorbitol (an osmotic stabilizer) improved cell viability [47]. Although we also observed oversized cells one day after the *top2* MC (Fig 2A and D), cell lysis was a rare event and sorbitol did not improve cell viability (Fig 2D-F). Therefore, we propose that the secondary ACD observed after the *top2* MC is the consequence of the steady decline of cell homeostasis resulting from loss of essential genes.

### On the long-term genetic consequences of the *top2* mitotic catastrophe

We have also compared the genomes of surviving diploids after 6h of Top2 inactivation. Even though we did not prove that all survivors came from a MC (i.e., they went through anaphase), the results shown in Figs 1-5, together with previous findings [13,24,25] lead us to conclude that most of these survivors likely came from a *top2* MC. Indeed, many of the observed chromosome variations and rearrangements can be best explained if cells go through anaphase in the absence of proper sister chromatid disjunction [11]. One interpretation of the strong bias for T-LOH events relative to I-LOH events (76 terminal and 3 interstitial) is that DSBs are repaired primarily through BIR (Fig. 5E) [11, 51]. In many previous studies, I-LOH represented a very significant fraction of the total LOH events. For example, in G_1_-synchronized cells treated with ultraviolet light, we observed a 1:3 ratio between T-LOH and I-LOH [40]. I-LOH requires both ends of the DSB to invade the other homolog during repair through homologous recombination, a condition difficult to fulfil in DSBs generated by cytokinesis [11].

According to a previous study [52], the frequency of DSBs in *top2* mutants is higher for long chromosome arms than short chromosome arms. Spell and Holm [50] explained this observation by suggesting that intertwinings on short chromosome arms could be resolved by passive diffusion off the end, whereas such intertwinings on long chromosome arms required Top2. Our results are also consistent with this interpretation. Chromosome IV and XII right arms are the longest in the yeast genome and are overrepresented in the T-LOH events (normalizing for the size of the arm). An unusual feature of the T-LOH data is the rDNA is under-represented as a breakpoint in the cXIIr T-LOH events. It was previously shown, using an assay that detects loss of inserted marker within the rDNA, that *top2* strains have substantially elevated rates of mitotic recombination in the rDNA [53]. One possible explanation of this discrepancy is that DSBs within the rDNA may be repaired by single-strand annealing between flanking copies of the rDNA genes [54], an event that would result in loss of an inserted marker without an interaction with the other homolog. Such an event could not be detected by the microarray analysis.

Another genetic alteration that is consistent with sister chromatid non-disjunction at the *top2* MC is the elevated presence of trisomies. A straightforward explanation for this is that the intertwining of sister chromatids in the *top2* strains often results in their co-segregation into one of the daughter cells (Fig. 5E). Although this type of non-disjunction would be expected to create equal numbers of monosomic and trisomic strains, it is possible that the monosomic strains have a competitive growth disadvantage and are, therefore, selected against. In *tel1 mec1* diploids, expected to enter anaphase with broken and/or underreplicated chromosomes, trisomies were five times more common than monosomies [55]. A similar bias towards trisomies was observed in *cdc14-1* diploids [51]; *cdc14* results in elevated levels of anaphase bridges because Cdc14 regulates condensin and Top2 actions in anaphase [56–58].

Lastly, the 100-fold enrichment in RUPD after the *top2* MC is also in agreement with models of genomic instability generated by sister chromatid non-disjunction. Two models are proposed for the generation of RUPD. One model involves two independent missegregation events occurring in successive divisions. In the other model, segregation of the chromosomes occurs in a manner similar to meiosis-I (Figs. S7 and S8). In wild-type cells, we demonstrated that the second model is correct [42]. From our data in the current study, we cannot distinguish between these models. The rate of a single non-disjunction event of chromosome V in a *top2* mutant is very high, about 3 x 10^-3^/division [23]. Thus, the likelihood of two independent non-disjunction events in a single division is (3 x 10^-3^)^2^ or 10^-5^ which is close to our observed rate of RUPD in the *top2-5* diploid. It is possible that both mechanisms contribute to the high rate of RUPD observed in the *top2-5* mutants. Another difference with our earlier study [42] is the low frequency (18%) of sectors reflecting RUPD and RCO recovered from the red/white Can^R^ sector assay. In our previous study, almost all sectored colonies reflected RCOs or RUPD events. It is likely that the high levels of aneuploidy observed in the *top2* mutants affects colony color by mechanisms unrelated to RCO or RUPD events on chromosome V.

In conclusion, we have characterized the consequences for the progeny of depleting yeast cells of Top2. We determined that the *top2* mitotic catastrophe leads to the sudden loss of the capability to divide again. Nevertheless, restrictions for cell division are not a consequence of immediate cell death as the progeny remain alive for several hours. In addition, survivors of the *top2* mitotic catastrophe carry genomic footprints that point towards sister chromatid non-disjunction and breakage of anaphase bridges as the source of the *top2*-driven genome instability. Overall, the *top2*-mediated mitotic catastrophe is highly deleterious for the cell progeny but it might also bring about highly unstable surviving clones.

## MATERIALS AND METHODS

### Strain construction

All strains used in this work are listed in Table S3. Gene deletions were achieved using standard PCR and transformation methods [25, 59]. To obtain the transformation products, genomic DNA of the corresponding strain in the Euroscarf yeast haploid *MAT***a** knockout collection was used as the PCR template. Primers used in the PCR bind 100-400 bps upstream and downstream the deleted gene ORF (Table S4).

The *top2-5* isogenic homozygous diploid derivative was generated through a one-step marker-free transformation approach that takes advantage of the α-factor hypersensitivity in the haploid *MAT***a** *bar1*Δ genotype (a proof of concept is provided in Fig. S9). Further details in supplemental information. The *top2-5* heterozygous diploids FM1873 and MD684 were obtained by crossing of haploid strain *top2-5* derivatives of PSL2 and PSL5 (see supplemental information). These two strains are isogenic with W303-1A and YJM789, respectively, and have been engineered to select and visually detect chromosome V rearrangements (Fig 5A) [38]. W303-1A and YJM789 differ by about 55,000 SNPs. Thus, these hybrid heterozygous diploids were also used for genome-scale detection of chromosome rearrangements by SNP arrays (see below).

Unless stated otherwise, all strains were grown overnight in air orbital incubators at 25 °C in rich YPD media (1% w/v of yeast extract, 2% w/v peptone and 2% w/v dextrose) before every experiment.

### Assays to assess clone survivability and capability of single cells to divide

A modified clonogenic assay performed on agar plates was used to assess survivability of the progeny after the mitotic catastrophe (details in supplemental information). The purpose of this assay was to determine if at least one of the resulting cells in the progeny was still able to raise a new cell population, irrespective of how many times the cell has divided and how many cells in the progeny are viable. Half-life values (t ½), the time in which the clone survival drops to 50%, were calculated adjusting the data to a four-parameter model using Graphpad Prism 7.

The time-lapse microcolony analyses were performed with a 40x long-range objective mounted on a Leica LMD6000 direct microscope (details in supplemental information). The cell density on the agar plate surface was set to 25 cells per 10,000 µm^2^. The cell volume of the original mother cell in all three frames was calculated assuming a perfect sphere and taking the cell diameter for calculations.

The single cell analysis by micromanipulation was performed on YPD plates with a Singer Sporeplay tetrad microdissector (details in supplemental information). Just 12 cells were harvested per plate in order to avoid prolonged incubations at 25° C at the beginning of the experiment.

The segregation and morphology of the histone-labelled nucleus (H2A-GFP) was analyzed by wide-field fluorescence videomicroscopy as described before [25]. Further details in supplemental information.

### Assays to assess cell vitality and cell death

All the colorimetric and fluorometric assays to assess metabolic competence, ROS production, and plasma membrane permeability were carried out in asynchronous logarithmic cultures grown overnight at 25 °C, adjusting the OD_600_ to 0.2 and incubated them for 24 h at 37 °C. Samples were taken at 0, 4 and 24 h and directly observed under the microscope. Fluorescence microscopy was used instead of flow cytometry because strains were already fluorescent for H2A-GFP. ROS were visualized with 10 μg/ml of 2′,7′-dichlorofluorescin diacetate (DCFH-DA; Sigma-Aldrich; D6883); and 3 μg/ml PI (Fluka; #81845) was used to count dead cells. Both dyes were directly mixed with a 200 μl aliquot of the culture and incubated 15 min at 37 °C in the dark. Cell bodies were considered ROS positive when the cytoplasm stained green in the absence of staining for PI. FUN® 1 (Invitrogen; F7030) staining was done washing 200 μl of each sample with water containing 2% D-(+)-glucose and 10 mM Na-HEPES, pH 7.2, resuspending the cells in the same buffer with 10 μM FUN® 1 and incubating 30 min at 37 °C in the dark. For MB (Sigma-Aldrich; M9140) staining, 1 µl of each sample was mixed with 1 µl of a 0.04% w/v MB solution in water onto a microscope slide and directly visualized (bright field). Staining with FITC-labeled annexin V was undertaken in spheroplasts as previously described [32]. Minor modifications to this protocol were employed; namely, we used the TACS Annexin V-FITC Apoptosis Detection Kit (R&D systems; 4830-01-K) as well as 60 units/ml of zymolyase (ZymoResearch; E1005) for cell was digestion.

### Analysis of genomic rearrangements using microarrays

Two similar protocols were used to expose heterozygous diploids to the *top2*-mediated MC. For both protocols, cells were grown in YPD to an optical density of 0.5-1. For one set of experiments (marked as E1 in the tables in the text), the cells were struck for single colonies on plates containing solid YPD medium, incubated for 6 hours at 37 °C, then incubated at room temperature until colonies had formed. For the second protocol, the cells grown at room temperature were harvested by centrifugation, and resuspended in 37 °C liquid medium, followed by incubation for 6 hours at 37 °C. Following this incubation, they were struck on YPD plates and incubated at room temperature until colonies were formed. For both protocols, the control cells were struck to room temperature YPD plates without incubating them at 37 °C. In experiments to detect sectored colonies, the YPD medium was replaced with solid omission medium lacking arginine and containing 120 µg/ml canavanine [39].

We detected LOH events, aneuploidy and UPD using SNP-specific microarrays similar to those used by [60]. The details of this procedure can be found in the supplemental information and have been described before [36, 37]. In analysis of single-colony isolates, whole-genome arrays were used (Gene Expression Omnibus [GEO] #GPL20144). For analysis of sectored colonies, we used microarrays specific for the right arm of V, and chromosomes I, III, and VIII (GEO #GPL21274).

## ABBREVIATIONS

MC: mitotic catastrophe
RCD: regulated cell death
ACD: accidental cell death
DSB: DNA double strand break
MB: methylene blue
PI: propidium iodide
ROS: reactive oxygen species
PS: phosphatidylserine
SNP: single nucleotide polymorphism
LOH: loss of heterozygosity
T-LOH: terminal LOH
I-LOH: interstitial LOH
UPD: uniparental disomy
RUPD: reciprocal UPD
rDNA: ribosomal RNA gene cluster

## AKNOWLEDGEMENTS

We thank other members of both labs and Kerry Bloom for fruitful discussions and help. We also thank Jessel Ayra-Plasencia, Nayra Cabrera-Quintero and Annika Lange for technical help in some of the experiments that required micromanipulation and microcolony counting. We thank Yang Sui for help with Figures S7 and S8.

