## Supplemental Text, Figures and Tables for "Cytological and genetic consequences for the progeny of a mitotic catastrophe provoked by Topoisomerase II deficiency"

**Interpretation of microcolony assays.**

Cells from asynchronous cultures were seeded on the surface of a plate at a cell density that optimized distance between cells and number of cells per field. Selected fields normally contain budded (S/$G_2$/M) and unbudded ($G_0$/$G_1$) cells, oscillating from a 1:1 to a 1:2 ratio. In our analyses, we concentrated on unbudded cells. A picture was taken before shifting the incubation temperature to 37 ºC (0h), then another picture of the same field was taken after the 37 ºC incubation (either 6h or 24h), and a final third picture was taken the day after, once the plate had been re-incubated at 25 ºC. The situation of each $G_0$/$G_1$ cell at the beginning of the assay was monitored after the 37 ºC incubations (6h or 24h) and after the plates were shifted back to 25 ºC. The landscape of possible outcomes is as follows (indicating "cell bodies" that may raise from the one-bodied $G_0$/$G_1$ cell):

| After 37 ºC (Top2 inactivation) | After 25 ºC reincubation (Top2 re-activation) |
|---|---|
| 0 (lysis) | --- |
| 1 (did not bud) | 0 (lysis) |
| | 1 (remained unbudded) |
| | 2 (able to bud once Top2 is back) |
| | 3, 4, 5, etc. (short-term budding capability) |
| | >20-50 (will raise a viable population) |
| 2 (did bud once without Top2) | 0 (double lysis) |
| | 1 (one body lysed; the other did not divide again) |
| | 2 (no more budding even after Top2 reactivation) |
| | 3 (one body able to bud once Top2 is back) |
| | 4, 5, etc. (both bodies able to bud*) |
| | >20-50 (at least one body/cell is viable) |
| 3 (did bud twice without Top2) | 0, 1, 2 (no more budding and some bodies lysed) |
| | 3 (no more budding even after Top2 reactivation) |
| | 4 (1 of 3 bodies rebudded once) |
| | 5 (2 of 3 bodies rebudded once*) |
| | 6, etc. (3 of 3 bodies rebudded once *) |
| | >20-50 (at least one body/cell is viable) |
| 4 (mother and daughter rebudded again*) | 0, 1, 2, 3 (no more budding and some bodies lysed) |
| | 4 (no more budding even after Top2 reactivation) |
| | 5, 6, 7, 8, 9 (1-4 of 4 bodies rebudded once*) |
| | >20-50 (at least one body/cell is viable) |
| Etc. | Etc. |
| * Other interpretations on the origin of these microcolonies are possible | |

Each $G_1/G_0$ cells could travel through any of the aforementioned categories. In the main text, we sometimes refer to their trajectory as, for example, 2 → 1. This indicates that one $G_1/G_0$ at 0h became 2 bodies after the 37 ºC incubation, but then one body lysed, and the other did not bud again despite reactivating Top2 (25 ºC). Likewise, 3 → m means that the $G_1/G_0$ cell became a triplet at 37 ºC and then formed a large microcolony after Top2 re-activation. Because the quantification of all trajectories is complex, as shown in the outcome landscape table, we opted for sunburst charts (Fig. 2B, 2F, 3E, 4A). The inner circle in the sunburst chart depicts proportions of cell bodies after the 37 ºC incubation. The outer circle depicts proportion of cell bodies after the final 25 ºC incubations for each situation observed in the inner circle. In order to aid with the interpretation of trajectories, we used different colors for different cell body numbers in the inner circle, but kept the colors of the inner circle for the same progeny in the outer circle. The changes in body numbers between the inner and the outer circles are indicated by numbers.

**Interpretation of SNP arrays to detect genome rearrangements.**

Each SNP array contains 25-base oligonucleotides that matched either the W303-1A-associated allele or the YJM789-associated allele for about 13,000 different SNPs (out of 55,000 SNPs). The selected SNPs are distributed evenly across the genome, giving an array resolution of ~1 Kb (yeast genome is 13 Mb, excluding the 1-2 Mb of the repetitive ribosomal DNA array). By measuring the relative amounts of hybridization to each oligonucleotide, we could detect loss of heterozygosity (LOH) (an event expected from mitotic recombination), as well as analyse deletions, duplications, and changes in chromosome number. Examples of these types of genome rearrangements are given in the plots shown Fig. 5E and S8. The Y-axis shows the normalized hybridization ratio to probes specific to the W303-1A form of the SNP (red) or the YJM789 form of the SNP (blue). Heterozygous SNPs have ratios of about 1; in LOH events, SNPs derived from one homolog have a ratio of near 2 and those derived from the other have a ratio near 0. One common pattern is a terminal LOH (T-LOH) event, which can reflect either a reciprocal crossover or a non-reciprocal type of recombination termed “break-induced replication” (BIR, Fig. 5E) [1]. A second type of LOH is an interstitial LOH event (I-LOH) in which a region of LOH is flanked by heterozygous. I-LOH events (gene conversions) result from the non-reciprocal transfer of DNA sequences between homologs. Two other classes of genomic rearrangements are a consequence of chromosome non-disjunction. Such non-disjunction events can result in trisomy or monosomy. An event in which one homolog is duplicated and another deleted is called “uniparental disomy” (UPD) and can reflect a non-disjunction event in which the two homologs segregate into different daughter cells.

### Extended Materials and Methods.

#### Construction of the *top2-5/top2-5* isogenic homozygous diploid (FM1730).

We employed the one-step marker-free transformation-based protocol described in Fig. S9. Briefly, the haploid *MAT***a** *bar1*Δ *top2-5 HTA2-GFP* strain used in most experiments (FM1386) was transformed with a PCR product obtained from a *MAT***α** haploid strain. This product is designed such that it can only recombine with the *MAT* locus but not with the silent *HML/HMR* loci. We counterselected against the *MAT***a** genotype by spreading 5 µg α-factor on the Petri dish surface before spreading the transformed cells. Colonies resistant to α-factor were collected after 3-4 days at 25 ºC and checked by PCR for the *MAT***a**, *MAT***α** or *MAT***a**/*MAT***α** genotypes (Fig. S9). *MAT***a**/*MAT***α** diploids were further confirmed by sporulation capability and 2N DNA content by flow cytometry [2].

#### Construction of the *top2-5/top2-5* hybrid heterozygous diploids (FM1873 and MD684).

The *top2-5* heterozygous diploids FM1873 and MD684 were obtained by crossing of haploid strain *top2-5* derivatives of PSL2 and PSL5. These two strains are isogenic with W303-1A and YJM789, respectively, and have been engineered to select and visually detect chromosome V rearrangements [3]. W303-1A and YJM789 differ by about 55,000 SNPs. The PSL2 *top2-5* (FM1830) and PSL5 *top2-5* (FM1832) haploids were constructed by transformation with a *top2-5:9xmyc:natNT2* product. This product was amplified by PCR from a CH326 strain derivative in which the *top2-5* allele had been tagged at 3' with sequences for 9 copies of the Myc epitope [4]. The heterozygous *top2-5/top2-5* diploid FM1873 was obtained by crossing FM1830 and FM1832. After realizing that FM1873 already carried genome alterations at 25 ºC (3-4 copies of cXIV and cXIIr T-LOH), other attempts to construct this diploid were undertaken. Previously, the FM1830 and FM1832 haploids were analysed and it was determined that FM1830 had 1-2 copies of cXIV. Thus, FM1830 was backcrossed with W303 to cure the strain of genome alterations. One spore (MD681) was identified that had the same genotype as FM1830 except that it had only one copy of chromosome XIV. This strain was crossed to FM1832 to generate the diploid MD684. Although these steps of construction were designed to generate a diploid that was isogenic with FM1873 lacking the aneuploidy of XIV and the T-LOH event on cXIIr, subsequent microarray analysis showed that MD684 still had three to four copies of XIV, although the T-LOH event on cXIIr was absent.

**Clonogenic survivability assays.**

Clonogenic assays were performed directly on agar plates to determine survivability of the progeny regardless the actual number of daughter cells originated at the restrictive condition. For this purpose, $10^2$ and $10^3$ cells (as estimated after counting cells in an asynchronous logarithmic culture with a Neubauer chamber) were spread onto a set of 14 YPD plates. These plates were then incubated for 0, 3, 6, 9, 12, 24 and 48 h at 37 ºC. After that, they were switched to 25 ºC to allow the growth of the survivors. Colonies were counted after 3-4 days and were normalized to the number of colonies grown without exposure to the restrictive temperature (0 h).

**Microcolony assays.**

For the microcolony analysis, ~1.5 x $10^5$ cells (counted by a Neubauer chamber) were spread onto a YPD (or YPD plus 1.2 M Sorbitol) plate to yield a density on the plate surface of around 25 cells per 10,000 $\mu m^2$. Defined positions on the plate were marked by piercing the surface with the needle of a Singer Sporeplay tetrad microdissector, using its 8 x 8 grid as a reference (12-16 fields in total). The plate was then transferred to a Leica LMD6000 direct microscope equipped with a 6.7x and 40x long-range objectives. The 6.7x was used to locate the marked fields and the 40x to take pictures of the cells in those fields (corresponding to 0 h, 25 ºC). Next, the plates were incubated at 37 ºC for 6 or 24 h before taking new pictures of the same fields. The procedure was repeated one more time after incubating the plate back at 25 ºC for 18-24 h. Finally, the same microcolonies in the corresponding three frames per field were identified by eye and categorized as indicated in the figure legends.

**Single cell analysis of the progeny by micromanipulation.**

For micromanipulation of the progeny the different strains were streaked on YPD Petri dishes. Unbudded cells were harvested with a Singer Sporeplay tetrad microdissector. Just 12 cells were harvested per plate and arrayed along the A file stage grid template in order to avoid prolonged incubations at 25º C. They were then incubated 6 h at 37 ºC and then observed under the microscope to count the number of cells that originated from the initial cell. Next, each cell that had at least a new bud was subjected to a mild attempt to separate the cell bodies using the needle and the vibration device. If successful, the largest body remained in the A file, whereas the other body was transferred to the B file. Finally, the plate was incubated 4 d at 25 ºC to search for survivors.

**Assays to determine segregation and morphology of the nucleus.**

The segregation and morphology of the histone-labelled nucleus (H2A-GFP) was analyzed by wide-field fluorescence videomicroscopy. An asynchronous culture was concentrated by centrifugation to $OD_{660}$ of 3 and spread onto YPD agar 90 mm Petri dishes. Patches were made from this plate and mounted on a microscope slide. They were incubated at 37 ºC for 24 h in high humidity chambers to avoid the patch to dry. The same fields were photomicrographed at 0, 6 and 24 h, or as reported before [5]. For each time point, a series of z-focal plane images (10 planes, 0.6 µm depth) were collected on a Leica DMI6000, using a 63x/1.30 immersion objective and an ultrasensitive DFC 350 digital camera, and processed with the AF6000 software (Leica).

**SNP microarrays.**

In brief, for most experiments, genomic DNA was obtained from single-colony isolates of experimental samples (incubated for six hours at 37 ºC) that was labeled with Cy5-dUTP. Two different types of microarray-control DNA were used and labelled with Cy3-dUTP. For some experiments, we used DNA purified from the FM1873 culture before exposure to 37 ºC. In other experiments, we used DNA from the *TOP2/TOP2* isogenic strain JSC24 [1]. Following labeling of the samples, the experimental and microarray-control DNA samples were mixed and hybridized to the microarrays [6]. The microarrays were then scanned at wavelengths 532 and 635 nm with a GenePix scanner, and analyzed by GenePix Pro software. Hybridization signals for Cy5 and Cy3 were normalized over the array to a value of 1. Additional steps of normalization are described in [6]. Following normalization, the ratio of hybridization of the experimental samples to the control samples for individual SNPs was 1 if the experimental strain was heterozygous.

### SUPPLEMENTAL FIGURES.

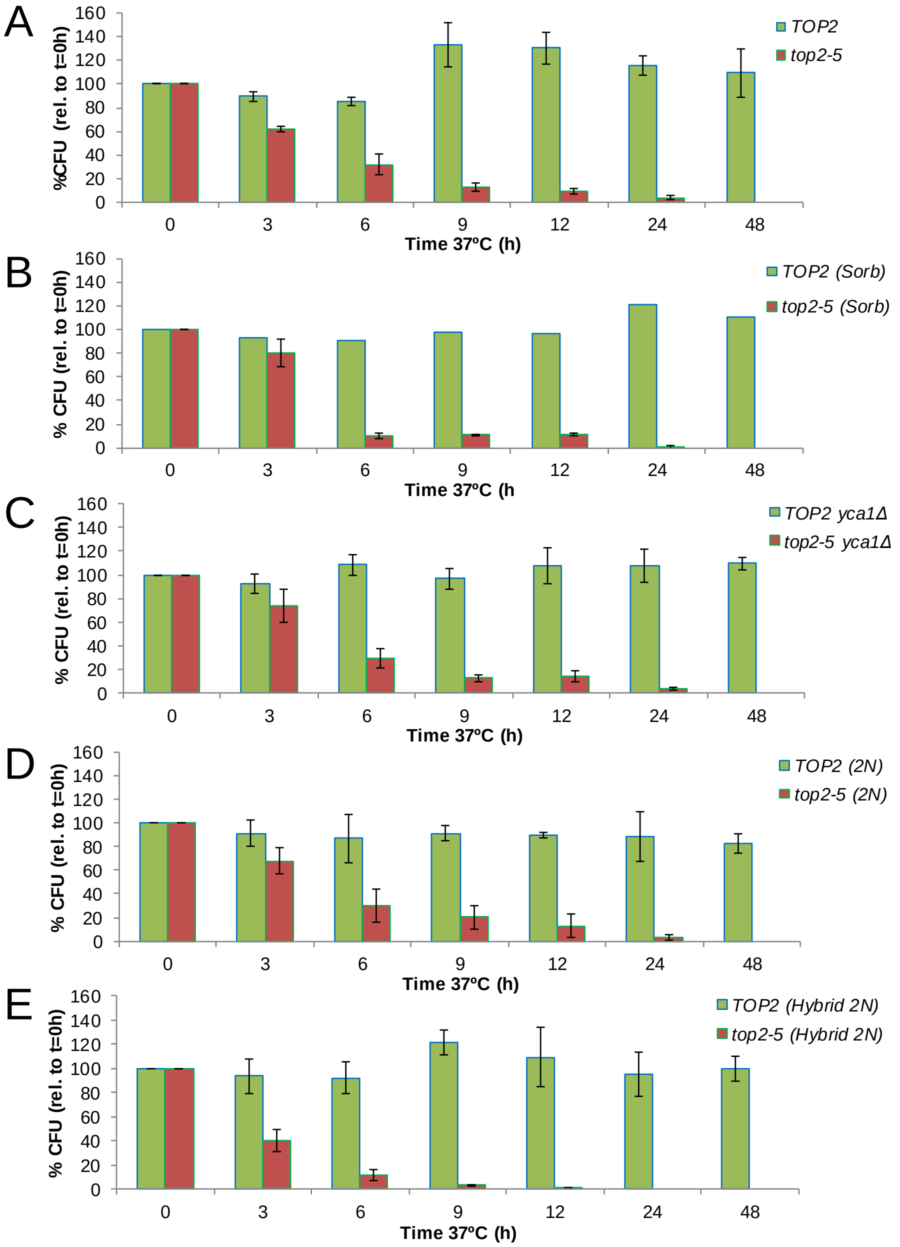


**Figure S1. Time course of clonogenic survivability of all *top2-5* strains and conditions studied and comparisons with their *TOP2* counterparts.** Asynchronous cultures growing at 25 ºC were spread onto several plates for each strain or condition. The plates were incubated at 37 ºC for different periods before transferring them 25 ºC. Four days after the initial plating, visible colonies (macrocolonies) were counted and normalized to a control plate which was never incubated at 37 ºC (0h). (**A**) haploid *top2-5* (FM1386) vs haploid *TOP2* (FM1419) on YPD. (**B**) The same two strains on YPD plus 1.2 M Sorbitol. (**C**) haploid *top2-5 yca1*Δ (FM1856) vs haploid *TOP2 yca1*Δ (FM1871) on YPD. (**D**) Homozygous diploid *top2-5* (FM1730) vs homozygous diploid *TOP2* (FM1732) on YPD. (**E**) Hybrid heterozygous diploid *top2-5* (FM1873) vs hybrid heterozygous diploid *TOP2* (FM2010) on YPD. These charts are related to Figures 1D, 2E, 3D, 4B and 5C, respectively.

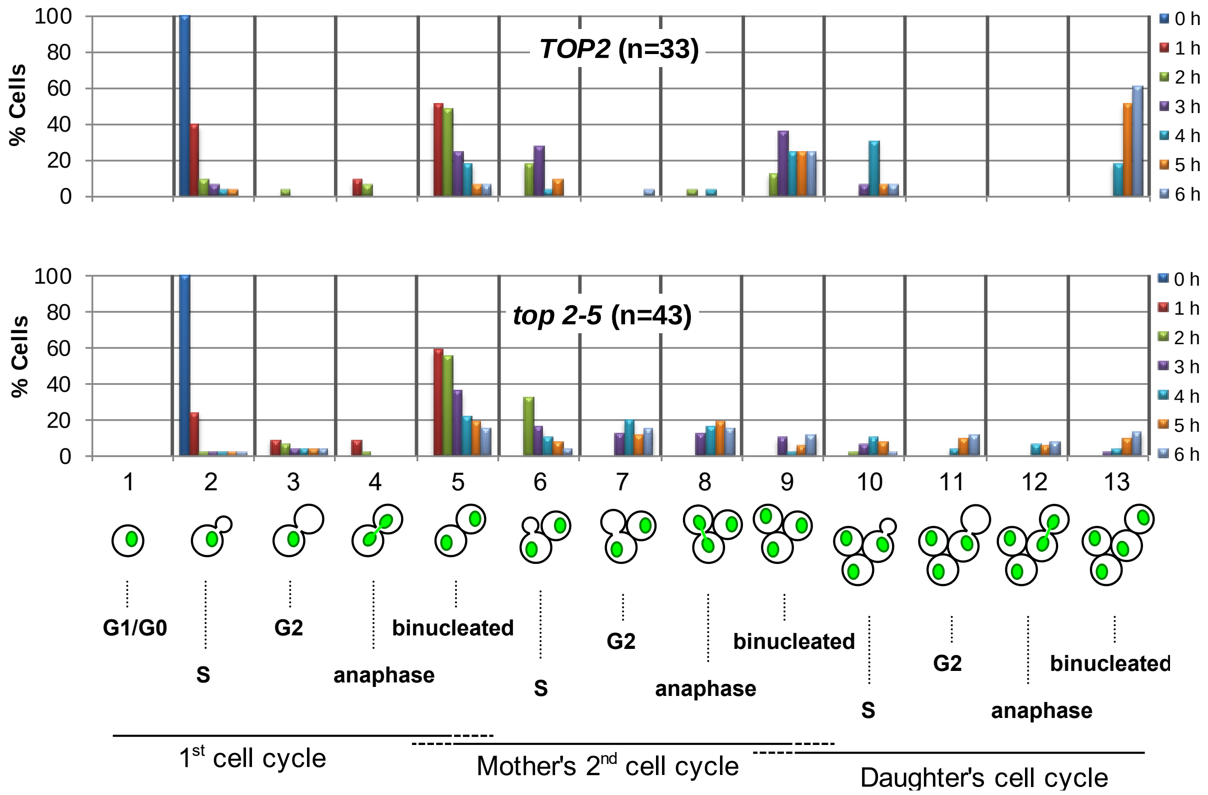


**Figure S2. Progression of cell division in budded *top2-5* cells at the time of the 37 ºC upshift.** HTA2-GFP cells carrying either the *TOP2* or the *top2-5* alleles were grown at 25 ºC concentrated to $OD_{600}$=3, spread onto YPD agarose patches and filmed under the microscope in a 37 ºC incubation. Total number of cells analysed (N) is indicated for each chart. Each hour, during a period of 6 h, a new photo was taken and each cell was moved to one of the indicated categories depending on whether it has budded, re-budded, segregated or attempted to split the nuclear mass, and whether any of these events have occurred in the mother or the daughter cell coming from the first division. The cells analysed in this experiment come from the same fields in which only unbudded cells at 0 h were followed in the Figure 2A in [5]. In this case, only budded cells at 0 h were considered.

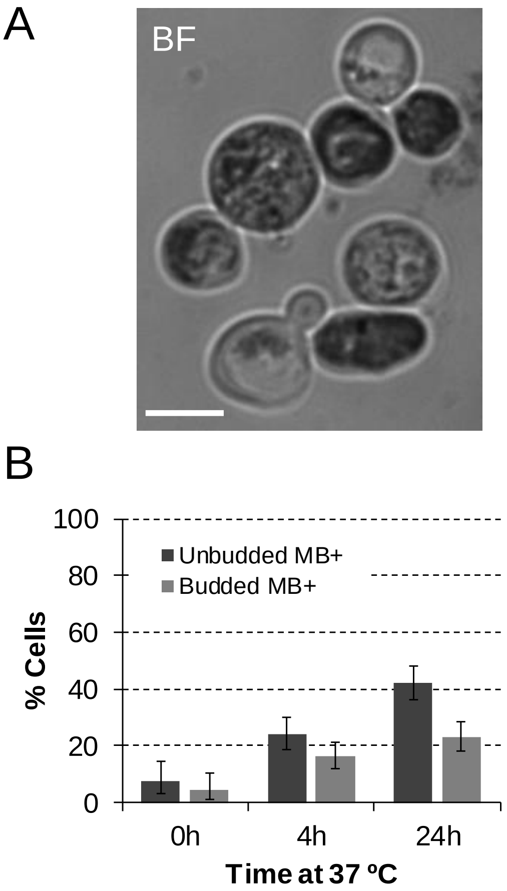


**Figure S3. Distribution of methylene blue staining in the *top2-5* population after incubation at 37 ºC for 24 h.** Related to Figure 3B. (**A**) Examples of cells that stained or not blue (dark grey in the image) after 24 h at 37º C. Note at the bottom of the image a triplet (mother with two attached daughters) where only one cell body (first daughter) has lost vitality. Scale bar depicts 5 µm; BF, bright field. (**B**) Percentage of MB+ cells at 0, 4 and 24 h at 37 ºC (± CI95). Cells were firstly classified as unbudded or budded (2-3 cell bodies). Budded cells were scored as MB+ if at least one body stained positive. The unbudded:budded ratio was ~2:1 in this experiment. Thus, the 40%:20% ratio after 24 h at 37 ºC implies that unbudded and budded cells are equally stained in relative terms.

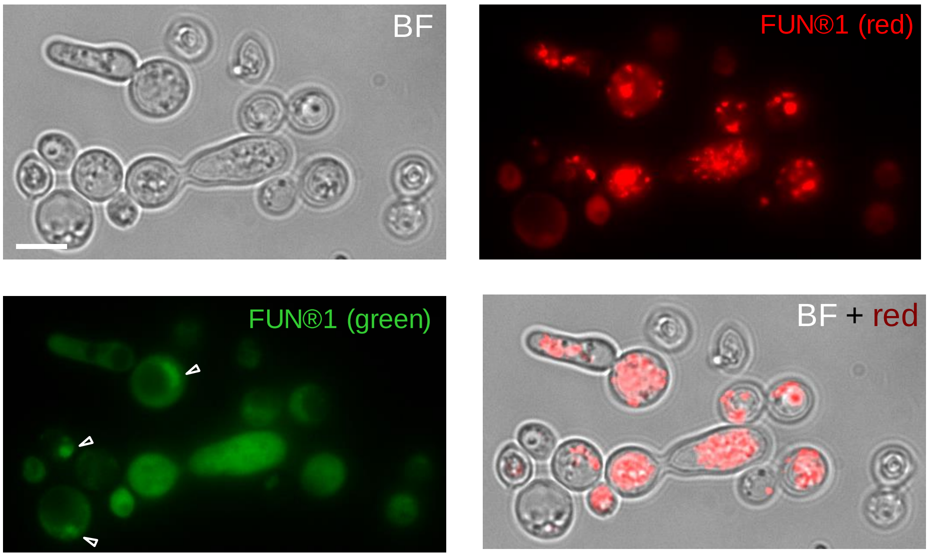


**Figure S4. Examples of *top2-5* cells stained with the vitality marker FUN® 1 after 24 h at 37 ºC.** Related to Figure 3C. In the green channel, hollow arrowheads point to the H2A-GFP signal. The cytoplasmic signal comes from unmetabolized FUN® 1. The brilliant aggregates in the red channel are metabolized FUN® 1 in cell bodies that retained a high vitality. Scale bar depicts 5 µm; BF, bright field.

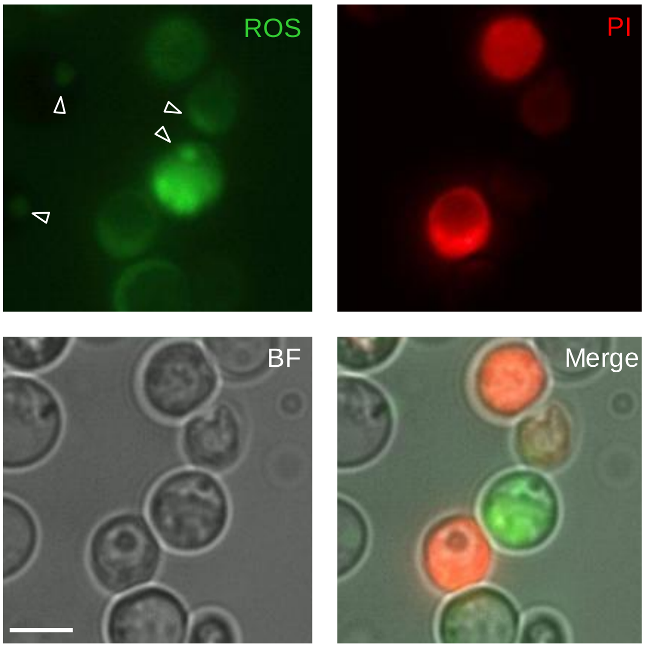


**Figure S5. Examples of *top2-5* cells co-stained for ROS and death after 24 h at 37 ºC.** Related to Figure 3C. In the green channel, hollow arrowheads point to the H2A-GFP signal. The cytoplasmic signal comes from ROS as reported with the DCFH-CA marker. The PI marker only stain cells that have lost plasma membrane impermeability (i.e., death cells). Note that cells are classified as ROS+ if they are bright green and PI negative. Scale bar depicts 5 µm; BF, bright field.

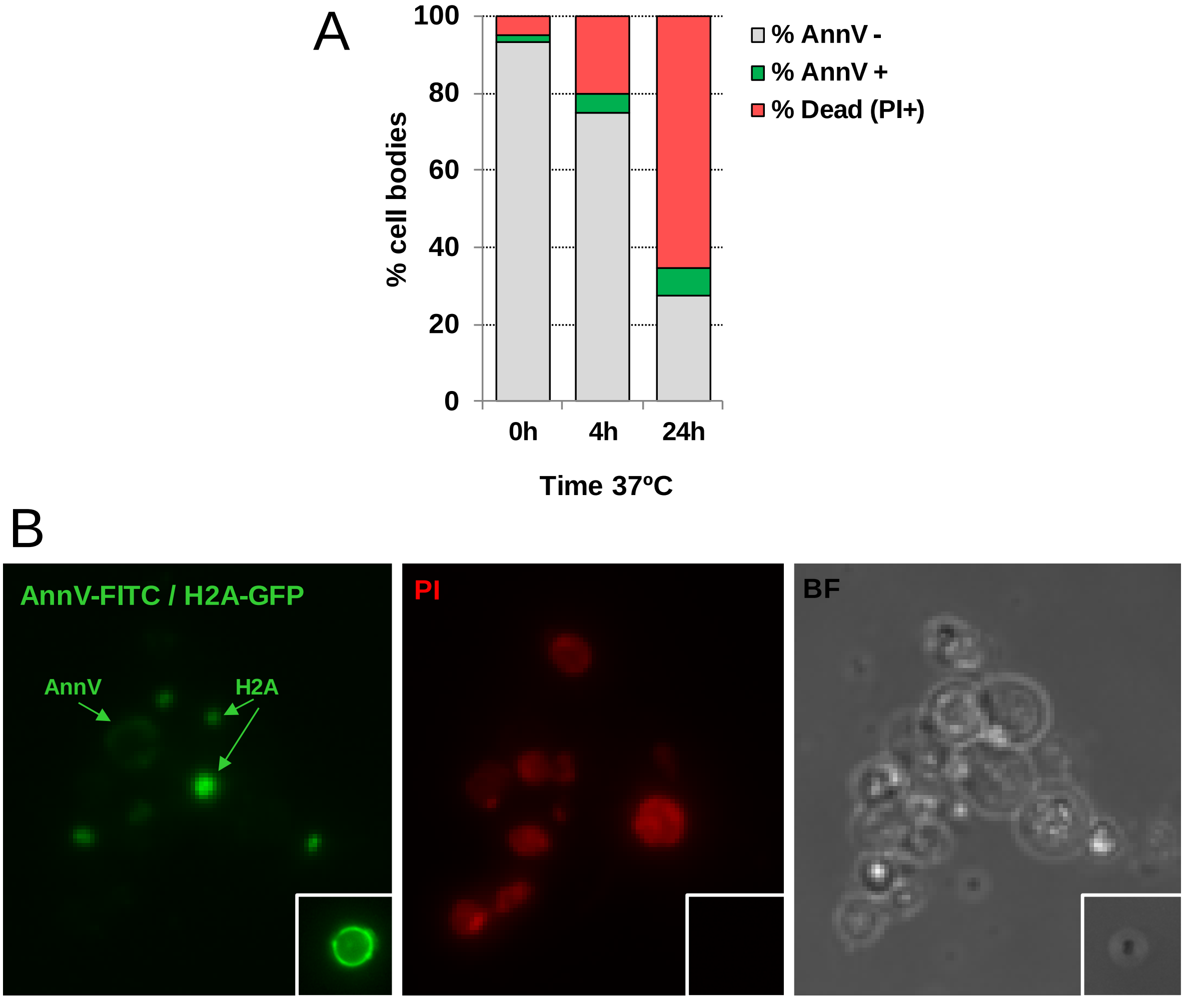


**Figure S6. Distribution of annexin V (annV) and propidium iodide (PI) co-staining in the *top2-5* population after incubation at 37 ºC.** (**A**) Percentage of cell bodies with different co-staining patterns at 0, 4 and 24 h at 37 ºC. (**B**) Examples of cell bodies with different co-staining patterns at 24 h. Note that cells also carry nuclear H2A labelled with GFP. Cell bodies were classified as AnnV+ when the fluorescent green signal was seen at the body periphery (as indicated). The inlet field depicts a cell membranous subparticle (probably an organelle) strongly stained by annexin V-FITC. These unusual examples (<1% of "bodies") were not considered in the quantification since a proper cell body was not inferred in the BF. Scale bar depicts 5 µm; BF, bright field.

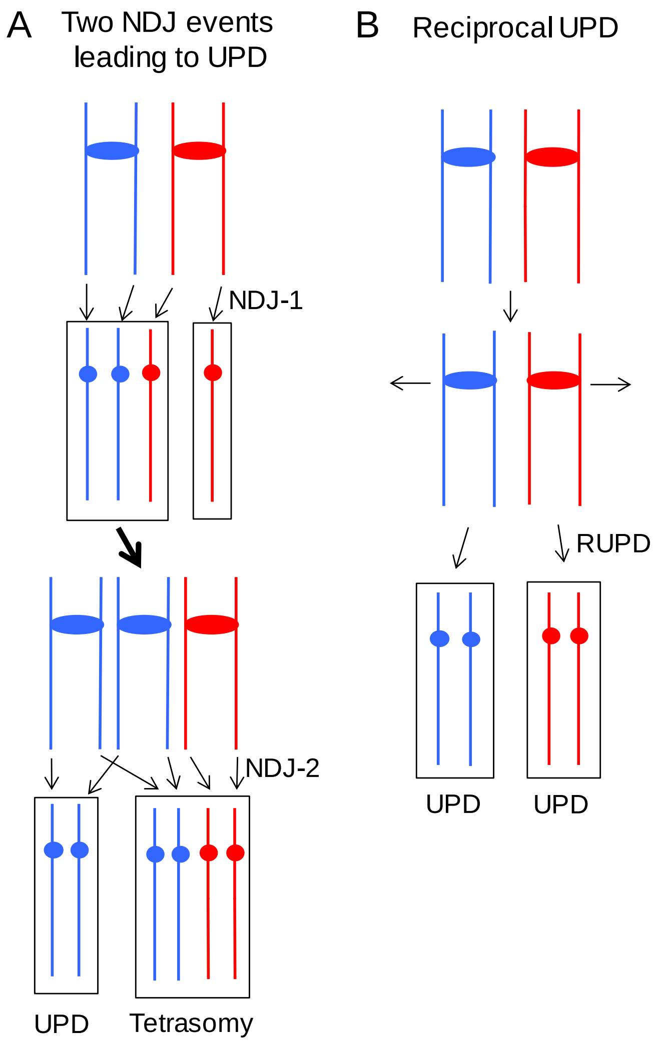


**Figure S7. Two pathways for generating UPD.** (**A**) Two non-disjunction (NDJ) events in different cell cycles can result in UPD. NDJ-1 results in one trisomic strain and one monosomic strain. If a second NDJ event occurs in the trisomic strain, a UPD isolate can be generated. (**B**) Reciprocal UPD. By this mechanism, two NDJ events in the same cell cycle results in two daughter cells with reciprocal patterns of UPD.

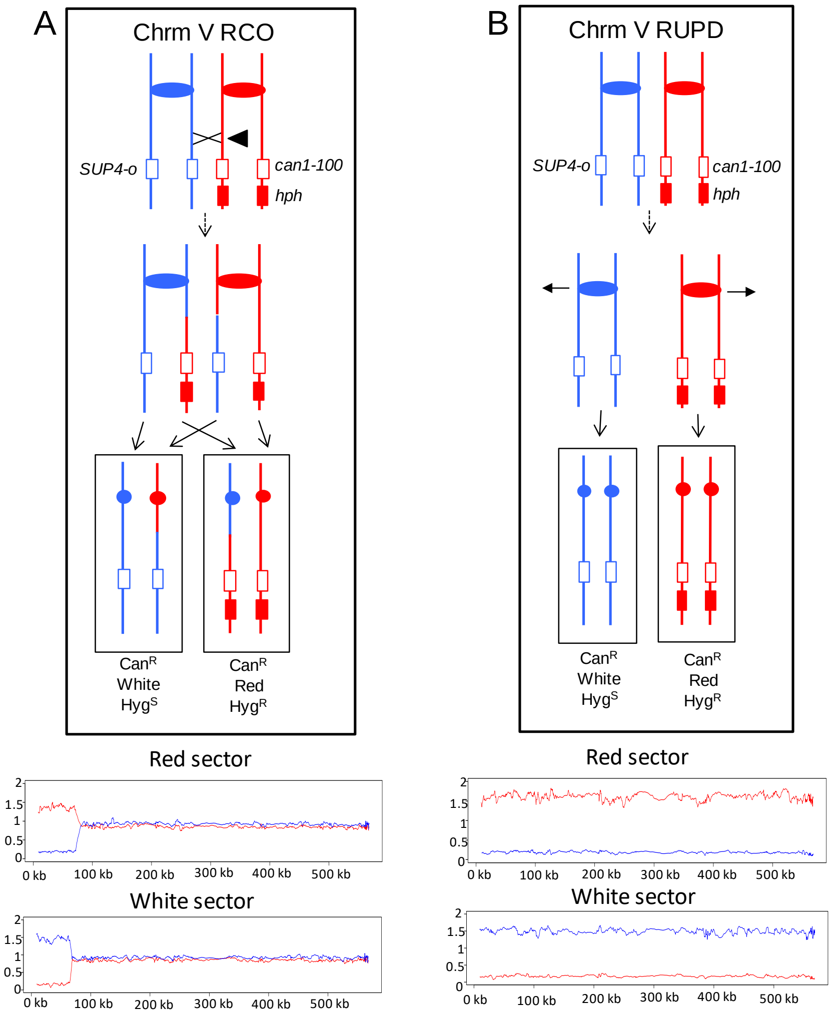


**Figure S8. Detection and analysis of reciprocal crossovers (RCO) and reciprocal UPD (RUPD).** The chromosome V homolog derived from YJM789 (shown in blue) has a *SUP4-o* gene (encoding an ochre suppressor) replacing the *CAN1* gene. On the W303-1A-derived homolog, chromosome V has the ochre-suppressible *can1-100* allele with the *hph* gene (resulting in hygromycin resistance) located centromere-distal to *can1-100*. In addition, the strain is homozygous for the ochre-suppressible *ade2-1* allele; diploids with this gene and 0, 1, or 2 copies of *SUP4-o* form red, pink or white colonies, respectively. Diploids without a genetic alteration form pink, canavanine-sensitive, hygromycin-resistant colonies. (**A**) A reciprocal crossover between the *can1-100* marker and *CEN5* can result in a red/white Can$^R$ sectored colony in which the white sector is sensitive to hygromycin. In the microarray pictures shown below the recombination event (FM1873-14 (E2) in Table S2), hybridization values for chromosome V from the red and white sectors are shown on the top and bottom panels, respectively, at the bottom of the figure. (**B**) A reciprocal UPD event on chromosome V would be expected to produce a red/white Can$^R$ sectored colony with the phenotypes identical to the RCO. The microarray patterns in each sector would be different. Microarrays derived from the red and white sectors of MD684.1.15 (E2) (Table S2) are shown in the top and bottom panels, respectively, at the bottom of the figure.

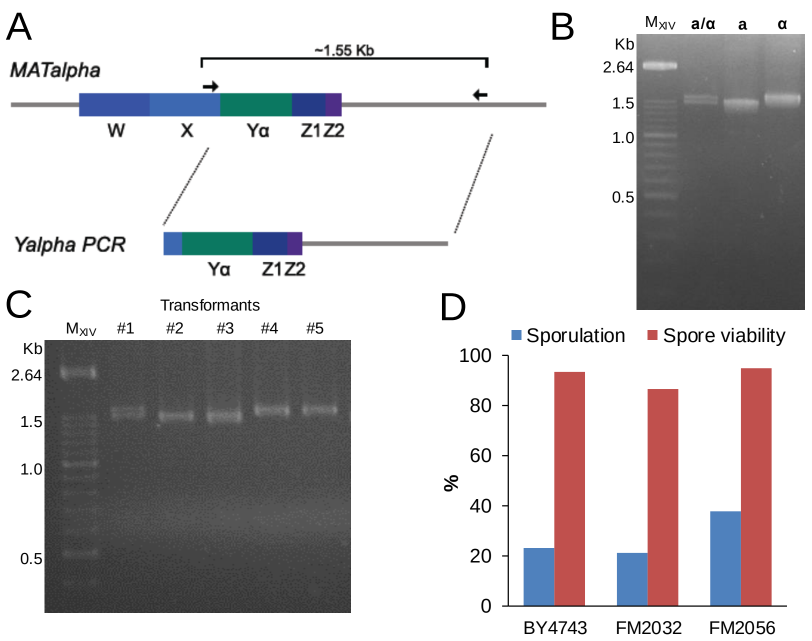


**Figure S9. One-step marker-free diploidization of *MAT*a *bar1Δ* strains. Proof of concept.** (**A**) Schematic of the *MATα* locus and the PCR-based strategy to obtain the *MAT***a**-to-*MATα* transformation product. The haploids *MAT***a** and *MATα* differ in the Y sequence within the *MAT* locus (chromosome III), but shared flanking sequences. The Y**α** is ~100 bps longer than Y**a** (747 bps vs 642 bps). The forward primer X-reg (Table S4) binds upstream of the Y region whereas the reverse primer MAT-R does so downstream, within the sequence adjacent to the MAT locus. Importantly, MAT-R does not bind downstream the silent *HML**α*** and *HMR***a** loci present in the same chromosome, which act during the mating type switching as template sequences for gene conversion at *MAT* [7]. (**B**) Gel separation of the three types of X-reg/MAT-R PCR products. The resulting PCR product is ~1.55 Kbps if the strain is a haploid *MATα*, whereas it is ~1.45 kb if the strain is a haploid *MAT***a** strain. A diploid would yield a double band that can be resolved in a 2% agarose gel. In these examples, the haploids and the diploid are the widely used S288C reference strains BY4741, BY4742 and BY4743. (**C**) Testing transformants for the one-step marker-free diploidization of *MAT***a** *bar1*Δ strains. Only *MAT***a** strains respond to the α-factor pheromone with a transient G1 arrest. The arrest is relieved by the Bar1 protease, which degrades α-factor extracelullarly. Knock-out mutants for *BAR1* become highly hypersensitive to α-factor; hence, in *bar1*Δ strains, α-factor can be used for the selection against the haploid *MAT***a** genotype. In this example, competent cells from the strain FM1932 (BY4741; *MAT***a** Δ*bar1::URA3*) were transformed with the Y**α** PCR product and spread onto YPD plates supplemented with 5 μg α-factor (spread on the plate surface). Five colonies were picked, re-struck onto YPD α-factor plates, and their genomic DNA tested for the genomic content at the MAT locus by X-reg/MAT-R PCR. Note that clone #1 is a diploid and clones #4 and #5 have become haploid *MATα*. Clones #2 and #3 were still haploid *MAT***a**. These strains probably come from spontaneous resistance to α-factor unrelated to genetic changes in the MAT locus (from our estimation: 0.2% over viable CFUs in a mock transformation experiment). (**D**) Sporulation efficiency and spore viability of the diploid obtained through the one-step marker-free method in comparison to diploids obtained through mating. BY4743 (*MAT***a**/α *BAR1*/*BAR1*); FM2032 (control *MAT***a**/α *bar1*Δ/*bar1*Δ obtained by mating the corresponding *bar1*Δ haploids); FM2056 (clone #1 from panel C).

**SUPPLEMENTAL TABLES.**

**Table S1. P-value of cross comparisons between the number of cell bodies generated after the 37 ºC for 6 h → 25 ºC for 16 h regime.** Comparison between microcolonies of ≥2 cell bodies after the 37 ºC incubation and re-budding (for at least one cell body) after the 25 ºC downshift were performed in 2x2 contingency tables using a one-tailed Fisher's exact test.

| Strain/condition | Microcolonies of ≥2 cell bodies at 37 ºC (N) | Microcolonies that rebudded after 25 ºC downshift (N) | P-value against *top2-5* (first row) |
|---|---|---|---|
| *top2-5* | 160 | 26 | - |
| *top2-5* (Sorb) | 89 | 21 | 0.1064 |
| *top2-5 yca1*Δ | 152 | 35 | 0.086 |
| *top2-5/top2-5* | 90 | 42 | < 0.0001 |

**Table S2. Summary of chromosome V events in R HygR/W HygS sectored colonies of FM1873 and MD684 strains.**

| Strain name[1] and sector pattern[2] | Genomic alterations[3] |
|---|---|
| FM1873-1 (E2) R $Hyg^R$/P-W $Hyg^S$ | UPD on V in “correct” direction in red sector; partial UPD in “incorrect” direction on V in white sector. |
| FM1873-2 (E2) R $Hyg^R$/W $Hyg^S$ | UPD on V in “correct” direction in red sector; T-LOH on V in “correct” direction in white sector. |
| FM1873-4 (E2) R $Hyg^R$/W $Hyg^S$ | RCO on V. Breakpoint at 134 kb in red sector and 144 kb in white sector. |
| FM1873-12 (E2) R $Hyg^R$/W $Hyg^S$ | RUPD on V. |
| FM1873-14 (E2) R $Hyg^R$/W $Hyg^S$ | RCO on V. Breakpoint at 67 kb in red sector and 76 kb in white sector. |
| FM1873-26 (E2) R $Hyg^R$/W $Hyg^S$ | UPD in correct direction on V in red sector; T-LOH event in correct direction in white sector. |
| FM1873-32 (E2) R $Hyg^R$/P $Hyg^S$ | RUPD on V |
| FM1873-35 (E2) R $Hyg^R$/P $Hyg^S$ | RUPD on V |
| FM1873-37 (E2) R $Hyg^R$/P $Hyg^S$ | RUPD on V |
| FM1873-41 (E2) R $Hyg^R$/P $Hyg^S$ | RUPD on V |
| FM1873-45 (E2) R $Hyg^R$/P $Hyg^S$ | UPD on V in the correct direction in the red sector (two copies of W303-1A-derived chromosome; white sector has one copy of YJM789-derived homolog and none of W303-1A-derived homolog |
| FM1873-49 (E2) R $Hyg^R$/P $Hyg^S$ | No obvious changes on V in red sector; white sector has UPD on V in correct direction. |

| | |
|---|---|
| FM1873-50 (E2) R $Hyg^R$/P $Hyg^S$ | RUPD on V |
| FM1873-54 (E2) R $Hyg^R$/P $Hyg^S$ | RUPD on V |
| FM1873-70 (E2) R $Hyg^R$/P $Hyg^S$ | RUPD on V |
| FM1873-85 (E2) R $Hyg^R$/P $Hyg^S$ | RUPD on V |
| FM1873-101 (E2) R $Hyg^R$/P $Hyg^S$ | RCO on V. Breakpoints at 95 kb and 86 kb in red sector. The red sector has event indicative of a G1-associated DSB, repaired in G2. Breakpoint at 93 kb in white sector. |
| FM1873-105 (E2) R $Hyg^R$/P $Hyg^S$ | RUPD on V. |
| FM1873-106 (E2) R $Hyg^R$/P $Hyg^S$ | RUPD on V. |
| FM1873-112 (E2) R $Hyg^R$/P $Hyg^S$ | RUPD on V. |
| FM1873-1 (C2) | UPD on V in “correct” direction in red sector; partial UPD in “incorrect” direction on V in white sector. |
| FM1873-2 (C2) | UPD on V in “correct” direction in red sector; no clear event on V in white sector. |
| FM1873-3 (C2) | No detectable events on V. |
| FM1873-7 (C2) | No detectable events on V in red sector. In white sector, V has terminal LOH (about 110 kb). |
| FM1873-14 (C2) | RUPD on V. |
| FM1873-19 (C2) | UPD on V in red sector; no obvious change on V in white sector. |
| FM1873-20 (C2) | RUPD on V. |
| MD684.1.15 (E2) | RUPD (V) |
| MD684.1.17 (E2) | RUPD (V) |

| | |
|---|---|
| MD684.1.49 (E2) | RUPD (V) |
| MD684.1.61 (E2) | Red sector looked like haploid strain (all homologs derived from W303-1A with elevated signal, all derived from YJM789 with reduced signal. UPD on V in white sector |
| MD684.1.65 (E2) | Red sector looked like haploid strain (all homologs derived from W303-1A with elevated signal, all derived from YJM789 with reduced signal). In white sector: UPD on V |
| MD684.1.73 (E2) | Red sector looked like haploid strain (all homologs derived from W303-1A with elevated signal, all derived from YJM789 with reduced signal). In white sector: UPD on V |
| MD684.1.75 (E2) | Red sector looked like haploid strain (all homologs derived from W303-1A with elevated signal, all derived from YJM789 with reduced signal). In white sector, UPD on V. |
| MD684.1.83 (E2) | RUPD (V) |
| MD684.1.88 (E2) | Red sector looked like haploid strain (all homologs derived from W303-1A with elevated signal, all derived from YJM789 with reduced signal). In white sector, T-LOH on V (breakpoint at 56 kb). |

[1] Parentheses after the strain name indicate whether the strain was experimental (E2, incubated for six hours at 37 ºC in liquid) or control (C2, not incubated at the restrictive temperature).

[2] Strains either treated at 37 ºC for six hours (E) or untreated (C) at the restrictive temperature were plated on solid medium containing canavanine. After colonies were formed, we purified cells derived from red and white sectors, and determined whether cells from these sectors were $Hyg^R$ or $Hyg^S$ (as indicated in the table). The *hph* marker was located distal to *can1-100* on the W303-1A-derived chromosome. Thus, a reciprocal crossover (RCO) or reciprocal UPD event would be expected to produce a hygromycin-resistant red sector, and a hygromycin-sensitive white sector. Code: T-LOH (terminal LOH event), I-LOH (interstitial LOH event), Tri (trisomy), UPD (uniparental disomy), RCO (reciprocal crossover), and RUPD (reciprocal uniparental disomy).

[3] The arrays for sectored colonies were done with arrays that had dense SNPs for chromosomes I, III, V and VIII, but few SNPs on other chromosomes. Thus, we tabulate only events involving chromosome V.

**Table S3. Strains used in this work.**

| Strain name | Relevant genotype[a] | Origin |
|---|---|---|
| CH326 | (S288C) *MAT***a** *ura3-52 his4-539am lys2-801am SUC2+ top2-5* | D. Botstein[b] |
| CH335 | (S288C) *MAT***a** *ura3-52 his4-539am lys2-801am SUC2+ TOP2* | D. Botstein[b] |
| FM1386 | CH326; *H2A2(YBL003c):GFP:BleMX*; Δ*bar1::URA3* | F. Machin[c] |
| FM1419[e] | CH335; *H2A2(YBL003c):GFP:BleMX*; Δ*bar1::URA3* | F. Machin[c] |
| FM1856 | FM1386; Δ*yca1::kanMX4* | This work |
| FM1871[e] | FM1419; Δ*yca1::kanMX4* | This work |
| FM1730[f] | *MAT***a/α** *top2-5/top2-5* homozygous diploid (from FM1386) | This work |
| FM1732[e,f] | *MAT***a/α** *TOP2/TOP2* homozygous diploid (from FM1419) | This work |
| PSL2 | (W303a) *MAT***a** *ade2-1 can1-100 his3-11,15 ura3-1 trp1-1 V9229::HYG V261553::LEU2 RAD5* | T. Petes[d] |
| PSL5 | (YJM789) *MAT***α** *ade2-1 ura3 can1Δ::SUP4-o gal2 ho::hisG* | T. Petes[d] |
| FM1830[g] | PSL2; *top2-5:9myc:natMX* (1-2 x cXIV) | This work |
| FM1832 | PSL5; *top2-5:9myc:natMX* | This work |
| FM1873[g] | (FM1830 x FM1832) *MAT***a/α** *top2-5/top2-5* hybrid diploid (3-4 x cXIV, cXIIr t-LOH) | This work |
| FM2010 | (PSL2 x PSL5) *MAT***a/α** *TOP2/TOP2* hybrid diploid | This work |
| MD681 | PSL2 *top2-5:9myc:natMX* (FM1830 backcrossed with W303 to have 1 x cXIV) | This work |
| MD684 | (MD681 x FM1832) *MAT***a/α** *top2-5/top2-5* hybrid diploid (3-4 x cXIV) | This work |
| BY4743 | *MAT***a/α** *his3Δ1/his3Δ1 leu2Δ0/leu2Δ0 met15Δ0/MET15 LYS2/lys2Δ0 ura3Δ0/ura3Δ0* | Euroscarf collection |

| | | |
|---|---|---|
| FM1932 | (BY4741) *MAT***a** *his3Δ1 leu2Δ0 met15Δ0 ura3Δ0*; *Δbar1::URA3* | This work |
| FM1982 | (BY4742) *MAT***a** *his3Δ1 leu2Δ0 lys2Δ0 ura3Δ0*; *Δbar1::URA3* | This work |
| FM2032 | (FM1932 x 1982) *MAT***a**/**α** *bar1Δ/bar1Δ* | This work |
| FM2056[f] | (clone #1 in Fig S9C) *MAT***a**/**α** *bar1Δ/bar1Δ* (from FM1932) | This work |

[a] Semicolons separate independent transformation events during strain construction. Intermediate strains are omitted.

[b] Described in [8]

[c] Described in [5]

[d] Described in [3]

[e] These strains were used as *TOP2* controls during clonogenic assays (n=3 independent experiments). In all cases, 100% viability was maintained after 0, 3, 6, 9, 12, 24 and 48 h incubations at 37 ºC.

[f] These homozygous diploids were made through the one-step marker-free transformation-based protocol described in Figure S9.

[g] These strains were shown by SNP and copy number arrays to carry the genome alteration shown between brackets. For instance, the hybrid heterozygous *top2-5/top2-5* diploid FM1873 carried two genome rearrangements when compared to its isogenic *TOP2/TOP2* counterpart: 3-4 copies of cXIV and a t-LOH at cXII right arm.

**Table S4. Primers used in this study.**

| Primer name | Purpose | Sequence (5' to 3') |
|---|---|---|
| Yca1-F (-359) | To amplify Δ*yca1::kanMX* from gDNA | CAATGCATTGGATCTTATTGGC |
| Yca1-R (+1709) | To amplify Δ*yca1::kanMX* from gDNA | GTCGAAACAAGAAGAGCAAAC |
| Bar1-F (-196) | To amplify Δ*bar1::URA3* from gDNA | GCCAGCTATTCTGAAACACACCAC |
| Bar1-R (+2316) | To amplify Δ*bar1::URA3* from gDNA | AACAGTCTTAGGGAAGTAACGAG |
| Top2-S3 | To tag *TOP2* at 3' with *9xmyc:natMX* | GGAAAACCAAGGATCAGATGTTTCGTTCA ATGAAGAGGATCGTACGCTGCAGGTCGAC |
| Top2-S2 | To tag *TOP2* at 3' with *9xmyc:natMX* | TATAAAAAGAATGGCGCTTTCTCGGATAA ATATTATTCAATCGATGAATTCGAGCTCG |
| Top2-F (-175) | To amplify top2-5*:9xmyc:natMX* from gDNA | AAGACGCGCCAGTAGGACGC |
| Top2-R (+4511) | To amplify top2-5*:9xmyc:natMX* from gDNA | CGCACGATGTTTTTCGCCCAGG |
| Xreg-F | To amplify Yα region in the MAT locus (Yα transformation product) | TTGTTGGCCCTAGATAAGAA |
| MAT-R (+2894) | To amplify the MAT locus (Yα transformation product) | CAAGGGAGAGAAGACTTGTG |

### SUPPLEMENTAL REFERENCES.


1. Yin Y, Petes TD. Genome-Wide High-Resolution Mapping of UV-Induced Mitotic Recombination Events in Saccharomyces cerevisiae. PLoS Genet. 2013; 9.
